# Unveiling novel signaling roles for human KDELR3 and KDELR1

**DOI:** 10.1101/2025.03.17.643648

**Authors:** Federica Cecilia Palazzo, Yuta Amagai, Marco Dalla Torre, Xue Han, Tiziana Tempio, Matthias Feige, Jose Garcia Manteiga, Masaki Matsumoto, Michele Sallese, Kenji Inaba, Roberto Sitia, Tiziana Anelli

## Abstract

KDEL receptors (KDELRs) prevent the secretion of soluble chaperones and enzymes meant to reside in the endoplasmic reticulum. While a single KDELR exists in yeast (ERD2), three variants are present in mammals, displaying high sequence similarity (73-83%). However, the phylogenetic conservation of the differences and the diverse tissue distribution of the three KDELRs suggest functional specialization. Here we show that, while all three receptors can prevent the secretion of KDEL-bearing clients, KDELR1 and KDELR3 regulate the production of AGR2, a key assistant of mucin folding, in opposite ways. AGR2 transcripts increase dramatically upon silencing KDELR3 but decrease when KDELR1 is downregulated. Silencing ERp44, but not other ER residents, phenocopies KDELR3 knockdown, suggesting that AGR2 regulation depends on ERp44-KDELR3 interactions. Our findings identify a novel regulatory circuit that controls the molecular composition of the early secretory pathway based on specific interactions between KDELRs and ER residents.

## INTRODUCTION

In eukaryotes, most membrane and secretory proteins are translocated cotranslationally into the endoplasmic reticulum (ER), the first station of the early secretory pathway (ESP). During their journey along the ER, ER to Golgi intermediate compartment (ERGIC) and proximal Golgi stations, cargo proteins encounter a number of chaperones and enzymes that control their folding and ensure the quality of the products that are to be released or transported to their final destination. To optimize folding efficiency, the protein composition of ESP subregions -all connected by vesicular transport mechanisms-must be strictly controlled. The anatomy of the secretory pathway favors the stepwise maturation of complex secretory proteins. Thus, BiP and ERp44 act sequentially (in time and space) in the biogenesis of IgM polymers, favoring the assembly of µ2L2 subunits in the ER and their polymerization in downstream compartments (Anelli et al, 2007; Tempio et al, 2021 and references therein). Most ER folding assistants (e.g., BiP, PDI, CNX) are ubiquitous. In addition, tissue-specific ones exist that are essential for particularly demanding proteins, like Hsp47 for collagen (Ishida & Nagata, 2011), MZB/CNP5 for immunoglobulins (Ig) (Rosenbaum et al, 2014; Xiong et al, 2019), or AGR2 for mucins (Park et al, 2009).

In general, unlike outgoing cargoes, folding assistants do not leave cells and remain in the ESP. Sorting relies also on a C-terminal tetrapeptide present in most soluble ER residents (Munro & Pelham, 1987). The cognate yeast receptor, ERD2, is a seven-transmembrane domain protein primarily residing at the ERGIC and cis-Golgi regions, where it recognizes the HDEL motif in escaped soluble residents, retrieving them to the ER (Lewis et al, 1990; Pelham, 1989; Semenza et al, 1990). Three ERD2 paralogues are present in humans, called KDEL receptors (KDELRs), since KDEL (Lys-Asp-Glu--Leu) is the most common retrieval motif in mammals (Brauer et al, 2019; Hsu et al, 1992; Lewis & Pelham, 1992; Lewis et al, 1990; Townsley et al, 1993). The three isoforms (KDELR1, KDELR2, KDELR3) share high sequence similarity (Fig.1). Interestingly, however, the differences that characterize the three KDELRs are highly conserved in metazoans, suggesting functional specialization (Fig. S1). Along with the increased complexity of the secretome in mammals, also the tetrapeptide motifs have diversified into over two dozen variants (e.g., KDEL, RDEL, QSEL, KTEL, etc.) (Raykhel et al, 2007). Interestingly, the tetrapeptides found in different chaperones are often phylogenetically conserved (e.g., KDEL in BiP, RDEL in ERp44).

**Figure 1.**
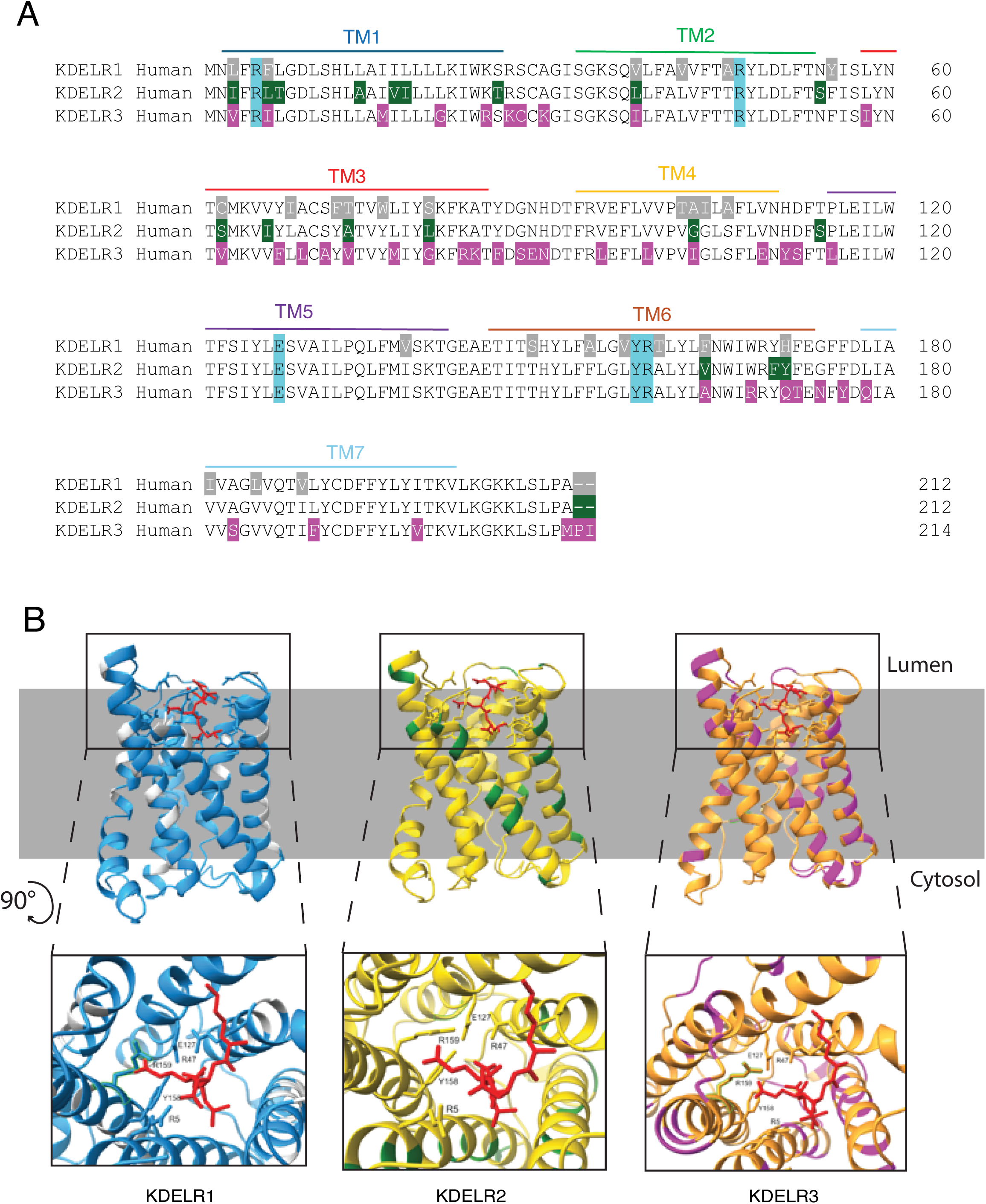
Sequence conservation amongst the three human KDELRs. A) Sequence alignment of the three human KDELRs. The amino acidic sequence of the three human KDELRs was aligned with Geneious Prime. The amino acids that differentiate each KDELR from the other two are in different colors: grey for KDELR1, green for KDELR2, and pink for KDELR3. In light blue are highlighted the conserved residues involved in the binding of the KDEL motif. The seven transmembrane (TM) regions are highlighted in different colors. B) Structural models of the three human KDELRs. The three-dimensional structures of the three human KDELRs were predicted using the AI tool Alphafold3. KDELR1 is depicted in light blue. In line with Fig. 1A, the amino acidic differences compared to the other KDELRs are highlighted in light grey. KDELR2 is in yellow, with the amino acidic differences highlighted in dark green. In orange is KDELR3, with the differences in magenta. A zoomed in image of the luminal part of each receptor is represented in the bottom part (squared) after rotation by 90 degrees. Bound KDEL peptides are represented in red, with the contacting residues in the KDELRs highlighted as sticks.

Does this complexity reflect redundancy or, rather, a functional specialization? The lack of antibodies efficiently discriminating the three KDELRs and the possible artifacts caused by their overexpression so far impeded a precise clarification of their function and localization (Cabrera et al, 2003; Capitani & Sallese, 2009; Townsley et al, 1993). However, the abundance and relative stoichiometry of the three KDELRs vary in different tissues (proteinatlas.org), suggesting specific functions in sorting and perhaps signaling. In fact, compelling evidence revealed that KDELR engagement can activate G-coupled proteins, impacting membrane trafficking, cytoskeletal rearrangements, and cell metabolism and growth (Cancino et al, 2014; Giannotta et al, 2012; Jia et al, 2024; Kamimura et al, 2015; Pulvirenti et al, 2008; Ruggiero et al, 2015; Ruggiero et al, 2018; Solis et al, 2017; Yamamoto et al, 2003; Yue et al, 2021; Zhang et al, 2021).

In this study, we investigated whether and how the three human KDELRs play different functional roles. We combine imaging, biochemical, and omic-based analyses which together demonstrate that while all three receptors have the potential to retrieve their clients, KDELR1 and KDELR3 play opposite roles in regulating the production of the ER-resident chaperone AGR2. Signals emanating upon encounter between the KDELRs and their ligands may allow cells to adapt the composition of the early secretory apparatus to the quality and quantity of the proteins in transit.

## RESULTS

### KDELRs are differentially involved in the retrieval of ER chaperones and enzymes

To unveil potential functional differences among the three human KDELRs, we first investigated the extracellular release of a subset of soluble ER residents upon knocking down (KD) each of the three KDELRs singularly or in various combinations (Fig. 2A, B). Silencing efficacy was ensured by RT-qPCR (Suppl. Fig. S2A). Anti-KDEL antibodies decorated four main bands in the lysates of HeLa cells. As expected, all were detectable in the culture medium upon combined KD of the three KDELRs (Fig. 2A, 123, lane 16). Interestingly, clear differences emerged when KDELRs were silenced singularly or in couples. Whilst KDELR2^KD^ and, to a lesser extent, KDELR1^KD^ were sufficient to induce detectable secretion of BiP, KDELR3 silencing had no discernible effects. Moreover, the simultaneous KD of KDELR1 and KDELR2 (KDELR12^KD^) was as efficient as the triple KD, suggesting that KDELR3 plays a minor role in preventing the secretion of ER-resident proteins in HeLa cells. To further investigate this issue, the same blots were decorated with antibodies specific for ERp44, a folding assistant that cycles between the ER and *cis*-Golgi and is retrieved by KDELRs binding through its RDEL tetrapeptide (Anelli et al, 2003; Anelli et al, 2007). In line with its rapid transport from the ER to the Golgi (Palazzo et al, 2022; Tempio et al, 2021), ERp44 was extensively secreted upon triple KDELR silencing, becoming almost undetectable in the lysates of KDELR123^KD^ cells (lane 8), as well as upon double KDELR1 and 2 KD (KDELR12^KD^, lane 5). Traces of ERp44 were detectable in the medium upon KDELR1^KD^ (lane 10), while a considerable amount of it leaked out upon KDELR2^KD^ (lane 11). By contrast, ERp44 was not secreted upon KDELR3^KD^ (lane 12).

**Figure 2.**
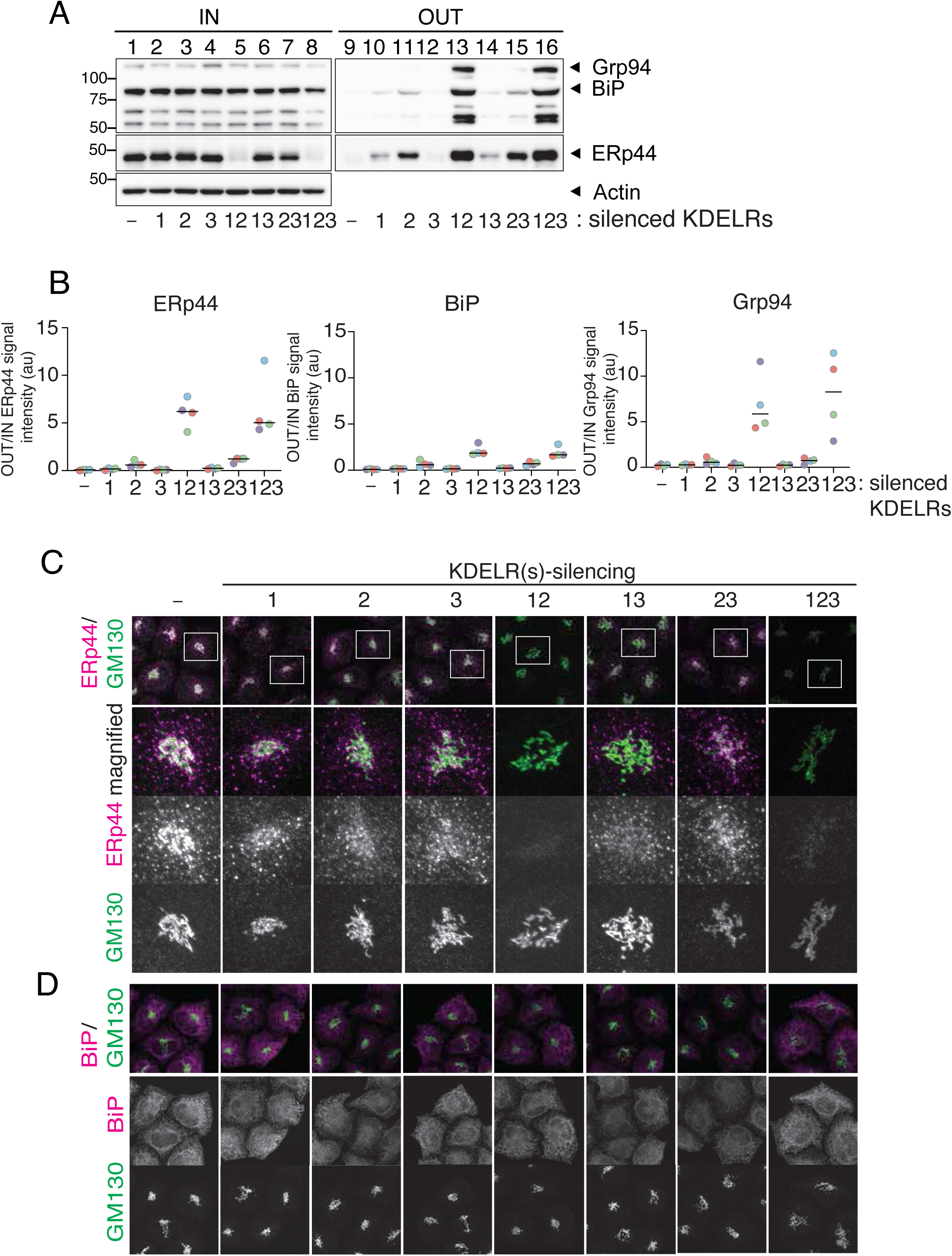
Role of KDELRs in the localization of soluble ER-resident chaperones. HeLa Kyoto cells were silenced with oligos specific for each human KDELR (alone or in combinations, as indicated), and secretion of ER-resident proteins monitored by WB (A-B) or imaging (C-D). A) *Secretion of KDEL-bearing proteins following KDELR downregulation.* The PVDF membrane was first decorated with anti-KDEL antibodies (upper panel). The same membrane was then washed and decorated with antibodies against ERp44. Actin served as loading controls. B) The signal intensities of ERp44, BiP, and Grp94 secreted by cells relative to those from whole cell lysates shown in (A) were quantified. Circles indicate the results from each experiment (N = 4 biological replicates). C) Immunostaining for ERp44 and GM130, a Golgi marker, in cells silenced for KDELR isoform(s) as indicated. Consistent with (A), the intracellular level of ERp44 was largely decreased upon double (KDELR12) or triple KDELR silencing (123). D) Cells silenced as in panel C were fixed and immunostained for BiP and GM130. Quantitative data for (C) and (D) are provided in Supplementary Figure S2B.

To gain further insight into the roles of KDELRs in ER protein retention, we analyzed the sub-cellular localization of BiP and ERp44. In the absence of KDELR2, ERp44 significantly accumulated in the Golgi (KDELR2^KD^ and KDELR23^KD^, Fig. 2C). Signals of intracellular ERp44 were almost undetectable upon double (KDELR12^KD^) and triple (KDELR123^KD^) silencing (Fig. 2C and Suppl. Fig. S2B), possibly reflecting its secretion into the medium (Fig. 2A, B). The rapid export of ERp44 from the ER towards the Golgi (Tempio & Anelli, 2020) likely makes the protein highly susceptible to KDELR downregulation. In this regard, ERp44 can be utilized as a useful indicator that sensitively reflects the cargo transport activities of KDELR1, 2, or 3.

To quantitatively analyze the Golgi localization of ERp44, Pearson’s correlation coefficients of ERp44 and GM130, a *cis*-Golgi marker protein, were calculated to describe the overlap/colocalization of their fluorescence signals (Amagai et al, 2023) (Fig. 2C, D and Suppl. Fig. 2C). Owing to its rapid exit from the ER, we expect that ERp44 would first accumulate in the Golgi and then progressively leave the cell upon RDEL deletion (Tempio et al, 2021) or inhibition of KDELR-dependent retrieval. Accordingly, Pearson’s correlation coefficient confirms that ERp44 is concentrated within the Golgi area upon KDELR2^KD^, indicating that KDELR2 is essential for efficient ERp44 retrieval. Quantitative analyses also demonstrated a reduced localization of ERp44 in the Golgi area (i.e., lower Pearson’s correlation coefficient of ERp44 and GM130) under KDELR1^KD^ and double KDELR13^KD^ compared to controls. These results suggest that KDELR1 can serve to retain ERp44.

In contrast to ERp44, BiP kept an ER-like distribution in all KDELR knockdowns (Fig. 2D and Suppl. Fig. S2B). BiP thus displayed a strong preference for localization in the ER, possibly because it interacts with other chaperones and a wide range of substrate proteins with the assistance of J proteins in this organelle (Cai et al, 2023; Meunier et al, 2002; Misselwitz et al, 1998). Like BiP, PDI is slowly secreted out upon deletion of its KDEL motif (Anelli et al, 2007) (see also Fig. 4).

Quantitative PCR data indicate that in the HeLa cells used herein, KDELR2 is more abundant than KDELR1, while KDELR3 is by far the least abundant of the three (Suppl. Fig. S2D). Thus, the primary role of KDELR2 in preventing the release of KDEL-bearing ER residents (Fig. 2) could reflect its greater abundance compared to KDELR1 or 3. To assess the efficiency of the three receptors in client retrieval, we expressed silencing-resistant forms of each KDELR in HeLa cells silenced for all three. As shown in Fig. 3, all isoforms, when over-expressed, were able to efficiently retrieve ERp44. In line with the overall similarity of their tetrapeptide binding regions (Fig. 1), proteomic analyses using the BioID2 protein-proximity labeling approach (Kim et al, 2016) revealed that the three receptors displayed largely overlapping interactomes when overexpressed in HeLa cells (Suppl. Fig. S3 and Table S1). The above data indicated that all three KDELRs could mediate efficient retrieval when overexpressed. Hence, their tissue-specific differential expression would strongly suggest functional specialization beyond cargo retrieval.

**Figure 3.**
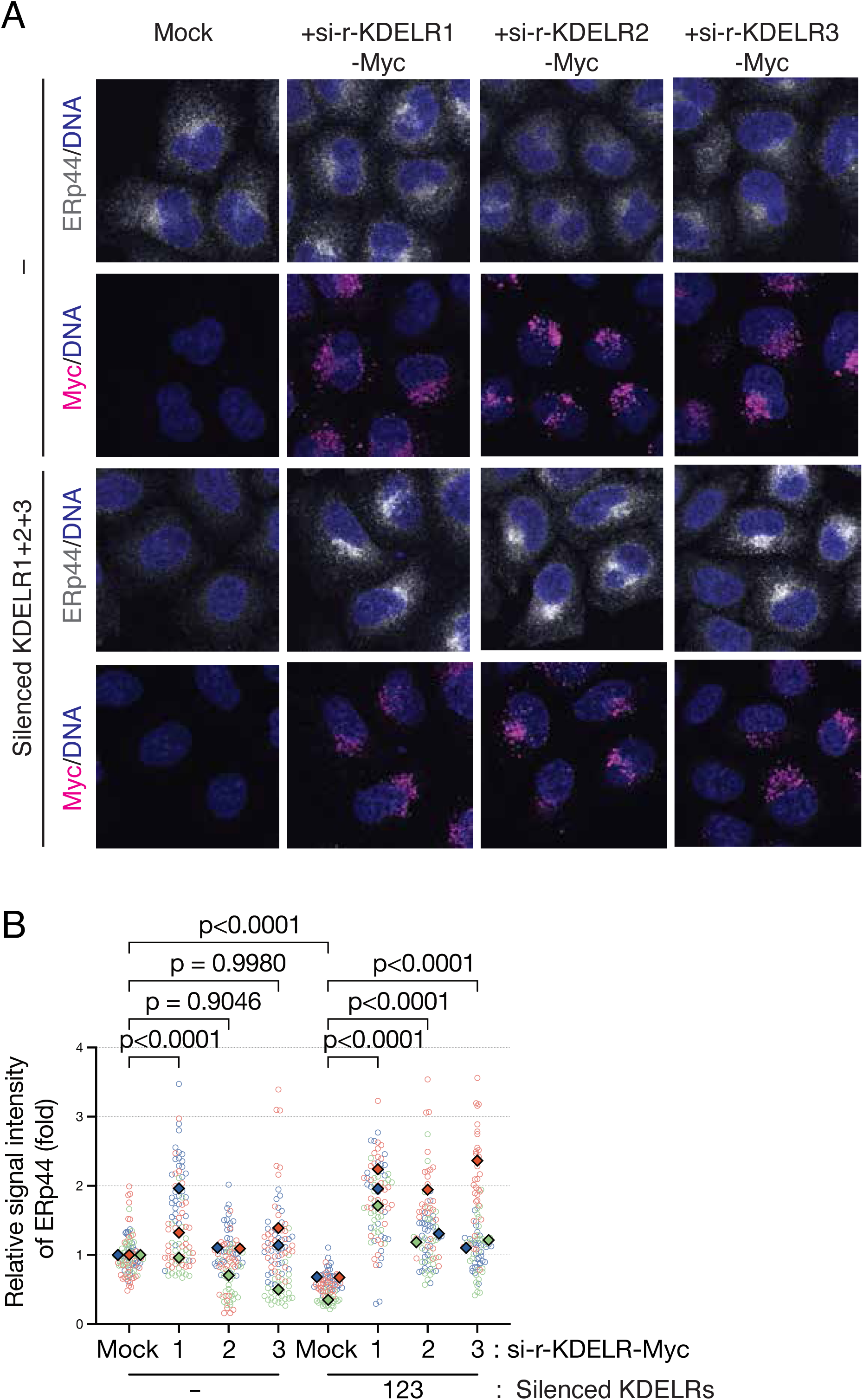
When overexpressed, all three KDELRs can prevent ERp44 secretion. A) HeLa Kyoto cells were transfected with plasmids expressing Myc-tagged siRNA-resistant-KDELR isoform (si-r-KDELR1/2/3-Myc). Cells were then transfected with siRNAs against KDELR1, 2, and 3 simultaneously or control reagents. After an additional 36 h incubation, cells were fixed and immunostained for ERp44 and Myc. Nuclei were stained with DAPI. B) The signal intensity of intracellular ERp44 shown in (A) was quantified and normalized relative to that of cells transfected with a mock plasmid and control siRNA. Circles indicate each data point, and rectangles indicate the median of each experiment (N = 3 biological replicates). One-way ANOVA followed by Tukey’s multiple comparison test was used for statistical analysis.

### Signals emanating from KDELRs regulate AGR2 production

Next, we analyzed the consequences of selective KDELR depletion on other ER resident proteins, including PDI, ERp46, and AGR2 (Fig. 4). A nitrocellulose membrane with the lysates and conditioned media of silenced and control cells was decorated with either anti-KDEL antibodies (as in Fig. 2A), or with specific antibodies against selected enzymes and chaperones. As expected, PDI, ERp46, and AGR2 are secreted upon triple KDELR123^KD^ (Fig. 4, lane 9). The identity of the two upper bands recognized by anti-KDEL (denoted X and Y) is unknown. Importantly, clear differences emerged when the distribution of the above KDELR clients was compared. ERp46 and X were almost undetectable in the lysates of cells lacking all three receptors (KDELR123^KD^, compare lanes 1 and 3 in Fig. 4A). As expected, ERp44 was extensively lost from triply silenced cells (see also Fig. 2A, lane 8). In contrast, PDI, Y, and AGR2 were almost as abundant in the lysates of KDELR123^KD^ cells as in mock-silenced cells (–), even though a fraction of them also accumulated in the medium. When either KDELR1 or KDELR3 was knocked down individually, none of the six proteins analyzed was secreted (lanes 10 and 12). In contrast, most of them partially leaked out upon KDELR2^KD^ (lane 11), confirming this isoform’s primary role in retrieving soluble ER residents in HeLa cells.

An unexpected role for KDELR3 emerged when AGR2 was analyzed (Fig. 4A). Decoration with specific antibodies revealed that abundant AGR2 accumulated intracellularly upon KDELR3^KD^ (Fig. 4A, lane 6). Surprisingly, a similar increase in AGR2 levels was observed upon ERp44^KD^ (Fig. 4A, lane 2). Detailed quantification of three or more similar experiments revealed that ERp44^KD^ favored AGR2 accumulation to a level similar to KDELR3^KD^ (Fig. 4B). Triple silencing had lower effects, likely because AGR2 was partially secreted out in the medium under these conditions (Fig. 4 A, B). ERp46, a PDI family member that is composed of three-thioredoxin-like domains followed by a C-terminal ER retention motif like ERp44 ^(Kojima^ ^et^ ^al, 2014)^ was rapidly secreted upon KDELR123^KD^ (Fig.4A). Importantly, however, its silencing did not significantly affect AGR2 levels (Fig. 4C), further suggesting a specific role of ERp44 in eliciting AGR2 accumulation. Considering that ERp44 and AGR2 share a similar CXXS active site, the increased AGR2 levels under ERp44^KD^ may reflect their compensatory functions. However, neither in HeLa nor in Caco-2 cells did AGR2^KD^ affect the protein levels of ERp44 (Fig. 4D). Thus, the levels of ERp44 specifically impact AGR2 accumulation.

**Figure 4.**
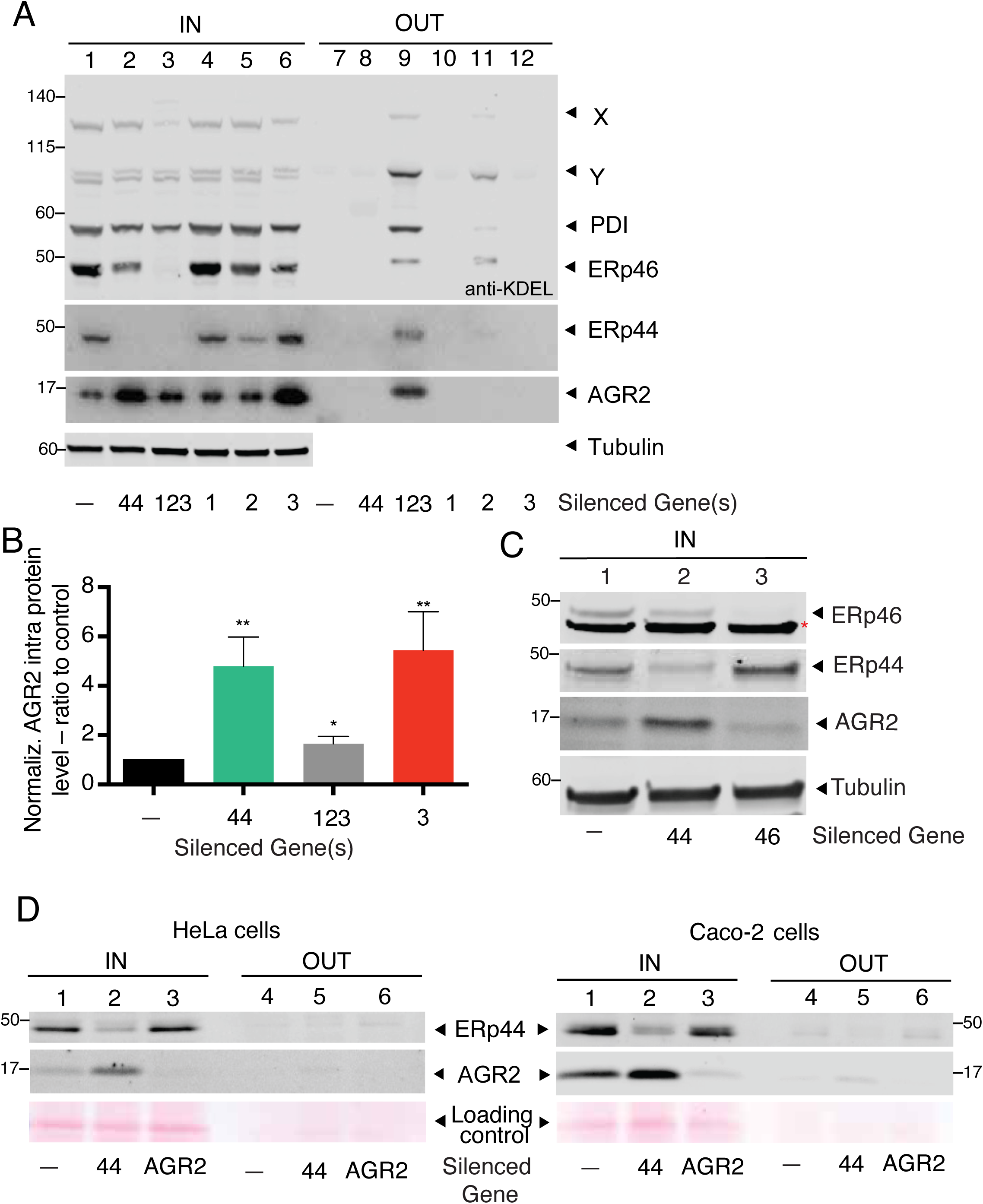
Different roles of human KDELRs. A) HeLa Milano or Caco-2 cells were silenced with KDELR-specific duplexes as indicated, and secretion of select ER resident chaperones was monitored by WB analyses. The nitrocellulose was first stained with anti-KDEL antibodies and subsequently with anti-ERp44 and anti-AGR2 as indicated. Tubulin was used as a loading control. The simultaneous KD of all KDELRs induced the extracellular release of almost all chaperones visualized in the WB. ERp44, ERp46, and the protein marked as X disappeared almost completely from cell lysates upon downregulation of all KDELRs, and partially under KDELR2^KD^. The intracellular levels of AGR2 increased upon downregulation of ERp44 or KDELR3. B) Densitometric quantification of WBs as the one shown in panel A. The bands decorated by the anti-AGR2 antibody were quantified densitometrically, and normalized against tubulin. The ratios obtained were normalized to untreated cells. C) HeLa Milano cells were treated with duplexes specific for silencing ERp44 or ERp46. Aliquots of their lysates were blotted and decorated with the indicated antibodies. Note that ERp46^KD^ did not impact AGR2 levels. D) AGR2^KD^ does not cause accumulation of ERp44, in HeLa Milano (left panel) or in mucin-producing Caco-2 cells (right panel). The ponceau signals provided loading controls.

### KDELR3 and KDELR1 control AGR2 at the pre-translational level

The above results prompted us to investigate whether transcriptional induction was responsible for AGR2 increase upon KDELR3^KD^ or ERp44^KD^. To this end, we performed RT-qPCR analyses (Fig. 5A). Clearly, the downregulation of KDELR3 (red bar) and ERp44 (green bar) induced a significant increase in AGR2 mRNAs. By contrast, KDELR1^KD^ (light blue bar) had a profoundly inhibitory role, leading to decreased AGR2 mRNAs. High induction levels were observed upon triple silencing (grey bar), implying that the inhibitory role of KDELR3 overrules the stimulatory one exerted by KDELR1. The levels of BiP and PDI mRNAs did not change significantly upon either KDELR3^KD^ or ERp44^KD^ (Fig. S5A), excluding a main role of the UPR in the emerging AGR2 regulatory circuits.

**Figure 5.**
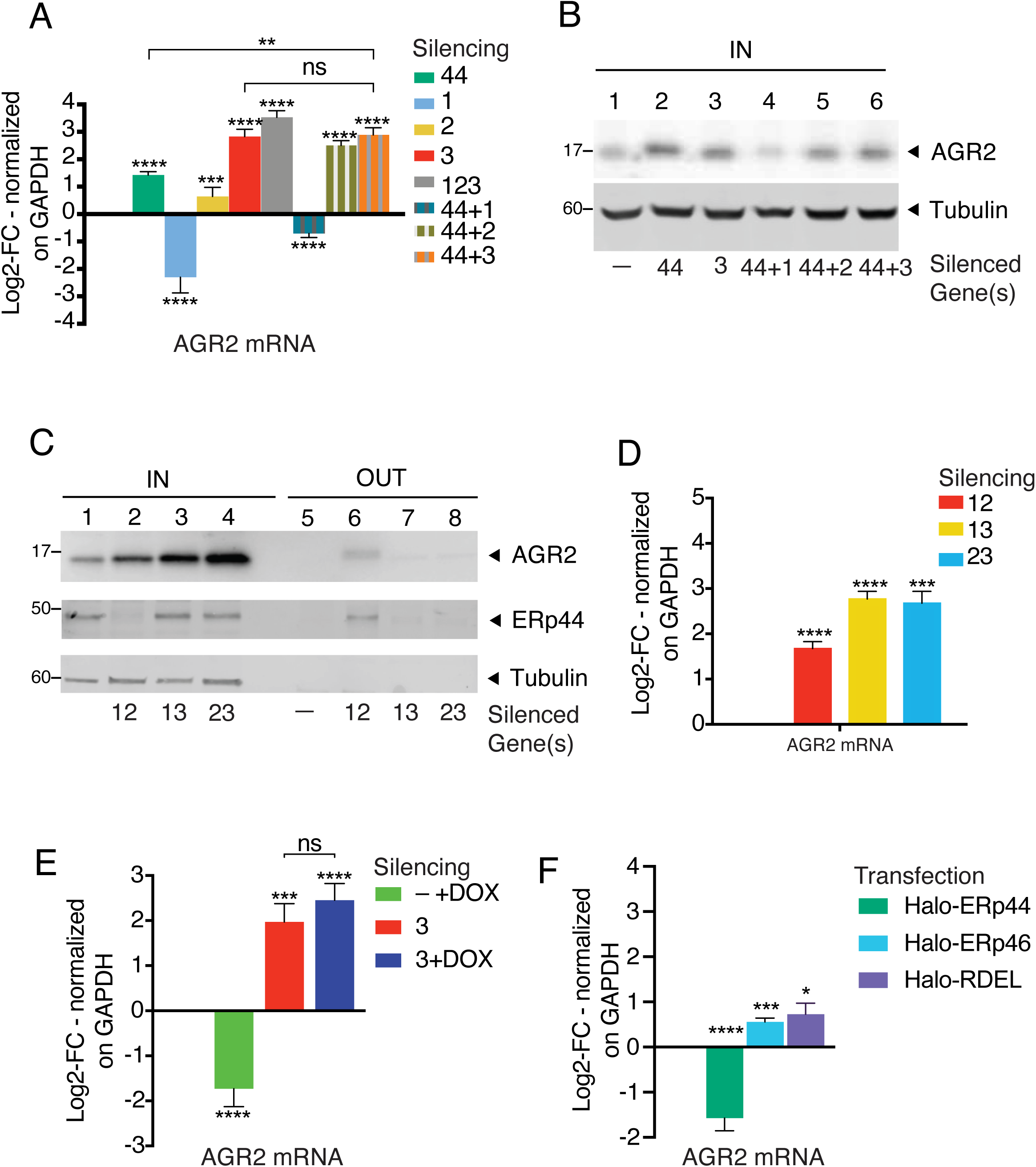
Opposite effects of KDELR1 and KDELR3 on AGR2. HeLa Milano cells were silenced as indicated and analyzed as described in Fig. 2. RT-qPCR analyses were performed to quantify mRNAs (panels A-D-E-F). Spent media were collected after 6 hours (panels B-C). For RT-qPCR analyses, GAPDH was used as a housekeeping gene to normalize the results obtained; data are expressed as Log2FC with respect to control conditions (that, by definition, corresponds to 0). *A)* AGR2 mRNAs increase upon ERp44^KD^ and/or KDELR3^KD^. 2 µg of RNA were used as starting material to perform RT-PCR and obtain the cDNAs used in the RTq-PCR analysis. Average of five independent experiments +/- SEM. *B)* Intracellular accumulation of AGR2. Tubulin is used as a loading control. *C)* AGR2 protein accumulates upon KDELR3^KD^ (also in combination with the KD of other KDELRs) and upon simultaneous silencing of KDELR1 and KDELR2, a condition in which most ERp44 is secreted. *D)* The panel shows the AGR2 transcript levels in cells treated as in C. mRNAs were quantified as in panel A. Average of five or more independent experiments +/- SEM. *E)* ERp44 rescues AGR2 inhibition in HeLa^ERp44KO^ cells via KDELR3. HeLa^ERp44KO^ cells, stably expressing doxycycline-inducible Halo-ERp44, were silenced for 72h for KDELR3 and then treated with 100µM doxycycline overnight. Transcripts were quantified as described in panels A and D. Average of three or more independent experiments +/- SEM. *F)* Neither HaloRDEL nor ERp46 activate KDELR3-dependent AGR2 inhibition. HeLa Milano ERp44 KO cells were transfected either with an empty vector or with Halo-ERp44, Halo-ERp46, and Halo-RDEL. After 48h, RNAs were extracted and quantified by RTq-PCR (see panel A). Average of three independent experiments +/- SEM.

To better elucidate the underlying mechanisms, we downregulated ERp44 in combination with each of the three KDELRs (Fig. 5A, B). The simultaneous downregulation of ERp44 and KDELR3 did not yield significant additional or synergistic effects, suggesting that the inhibitory role of ERp44 involves mainly KDELR3, and ERp44 and KDELR3 are epistatic in the same pathway. In contrast, simultaneous KDELR2 and ERp44 silencing (44+2, Fig. 5A, B) resulted in summative effects, suggesting that the two are not epistatic. Thus, the binding of ERp44 to KDELR3 yields signals that control the abundance of AGR2 transcripts, which are independent of ER stress (Fig. S5A). KDELR3^KD^ caused a significantly higher accumulation of AGR2 transcripts than ERp44^KD^ (Fig. 5A), suggesting that additional ligand(s), possibly clients of KDELR2, may influence AGR2 expression via KDELR3. These results indicate that KDELR3 inhibits, and KDELR1 stimulates, the accumulation of AGR2 transcripts and ERp44 is a key interactor of KDELR3.

Next, we analyzed the consequences of silencing two KDELRs simultaneously (Fig. 5C, D). Western blot (WB) analyses (Fig. 5C) confirmed that the least abundant of the three, KDELR3, played little role in retrieval, if any. However, KDELR3^KD^ had greater effects on the abundance of AGR2 protein and transcripts than double KDELR12^KD^. A significant rise in AGR2 (protein and mRNA) was evident upon dual KDELR13^KD^ silencing (Fig. 5C, D yellow bar), without extracellular accumulation of the protein. Upregulation of AGR2 was obtained also under KDELR23^KD^ (Fig. 5C, D light blue bar) when KDELR1 can boost AGR2 transcription. However, this increase was not followed by the leakage of the protein into the medium (Fig. 5C, lane 8), suggesting that KDELR1 alone can prevent AGR2 secretion. Surprisingly, AGR2 is induced (both at the protein and mRNA level) also under double KDELR12^KD^ (Fig. 5C, D red bar). This paradoxical finding can be explained by considering that ERp44 is rapidly secreted out of cells when KDELR1 and KDELR2 are silenced (see also Fig. 2). The few remaining molecules seem, hence, unable to unleash KDELR3-dependent inhibition.

Taken together, the above data reveal that in HeLa cells, KDELR2 is mainly responsible for client retrieval, whilst KDELR1 and KDELR3 also serve opposite signaling roles in regulating the abundance of AGR2.

### ERp44 acts via KDELR3 to regulate AGR2 expression levels

To confirm that ERp44 binding to KDELR3 activates signals that inhibit AGR2 expression, we used HeLa cells devoid of ERp44 (Giannone et al, 2022; Giannone et al, 2025) (HeLa^ERp44KO^) and reconstituted them with doxycycline (DOX)-inducible ERp44 (Fig. 5E). When these cells were treated with DOX to re-express ERp44, AGR2 decreased significantly (green bar). Importantly, ERp44 failed to dampen AGR2 transcription in cells lacking KDELR3 (compare blue and red bars), suggesting that specific interactions between KDELR3 and ERp44 generate epistatic signals that inhibit AGR2 expression. Accordingly, neither ERp46, a protein that, like ERp44, is rapidly lost by cells upon triple KDELR123^KD^, nor a neutral secretory reporter (spHalo-RDEL) (Tempio et al, 2021) rescued AGR2 inhibition when overexpressed in HeLa^ERp44KO^ cells. Thus, among the proteins over-expressed so far, only ERp44 was effective (Fig. 5F). This confirms that ERp44 plays an important role in inhibiting AGR2 production via KDELR3.

### Common transcriptional patterns emerge upon KDELR3 and ERp44 silencing

The above results indicated that specific interactions between select KDELRs and their clients generate signals controlling AGR2 abundance. To learn more about the pathways involved, we compared the transcriptomes of HeLa cells 72 hours after the silencing of each KDELR or ERp44. PCA analyses (Fig. 6A) revealed significant separation among the groups of samples, implying clear differences in their transcriptional patterns. As expected, AGR2 was amongst the most up-regulated genes in KDELR3^KD^ and ERp44^KD^ but down-regulated in KDELR1^KD^ (Suppl. Fig. S4A). Clear differences emerged when Gene Ontology pathways of modulated genes (False Discovery Rate – FDR-< 0.05 and log2FC > 1) were compared (Suppl. Fig. S4B). Silencing of any of the three isoforms severely affected pathways related to cell division, adhesion, and extracellular matrix, confirming previous results (Capitani & Sallese, 2009; Ruggiero et al, 2015; Ruggiero et al, 2018). KDELR2^KD^ and KDELR3^KD^ inhibited pathways specifically associated with mitotic chromosome segregation. ERp44^KD^ also inhibited genes related to cholesterol and lipid metabolism. Silencing either ERp44 or KDELR3 upregulated genes involved in G protein-coupled receptor signaling and BMP pathways. Moreover, downregulating KDELRs altered transcripts related to extracellular matrix and cell adhesion, again confirming previous results (Ruggiero et al, 2015; Ruggiero et al, 2018).

**Figure 6.**
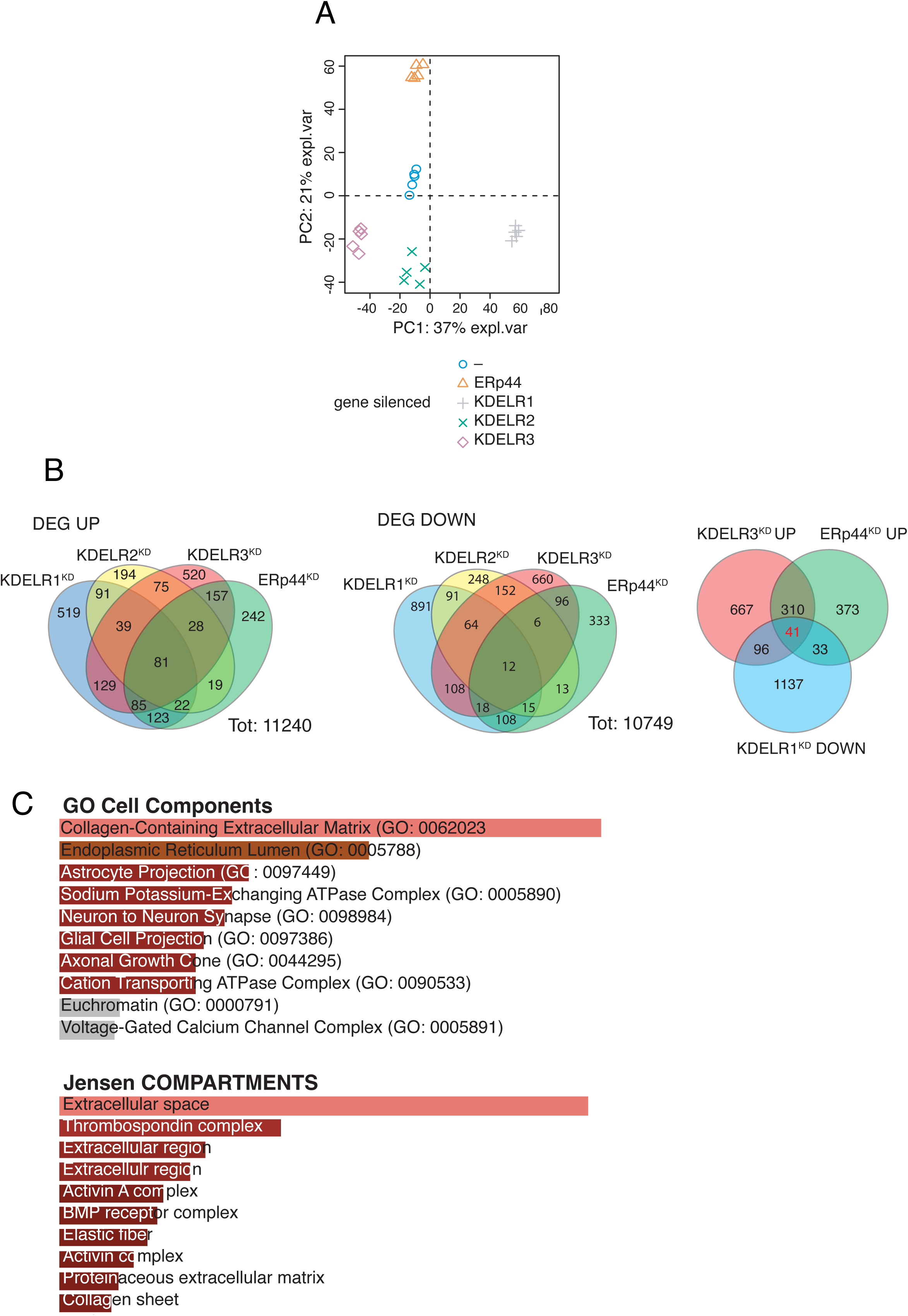
Transcriptional patterns of HeLa cells upon KDELRs downregulation. A) Principal Component Analysis Score plot PC1 vs PC2. The graph clearly shows the separation of the five experimental conditions. B) Overlap analysis of differentially expressed genes after silencing of KDEL receptors and ERp44. Left panel: genes up-regulated in the different conditions; central panel: genes down-regulated in the different comditions; right panel: genes up-regulated upon silencing of KDELR3 and ERp44 and dowregulated upon KDELR1, behavior observed for AGR2. C) Functional Enrichment using Enrichr of Gene Ontology Cellular Components and the Databases of Jensen Compartments of the 41 genes behaving like AGR2 showing enrichment in “Secreted Proteins” and “Endoplasmic Reticulum Lumen”.

Interestingly, other differentially expressed genes (DEGs) shared the AGR2 pattern: increased in KDELR3^KD^ or ERp44^KD^, decreased in KDELR1^KD^ (Fig. 6B). Gene Ontology analyses revealed these genes to be related to extracellular matrix composition (Fig. 6C). When data were filtered based on statistical significance only (FDR < 0.05), some genes related to the ESP and glycosylation emerged with the AGR2 trend (Suppl. Fig. S5B). Among these, ST6GAL1, MAN1A1, FUT8 (genes involved in glycosylation), ERp29 (an ER chaperone), and CD82 (a gene previously demonstrated to be partially dependent on KDELR3 (Marie et al, 2020; Zhu et al, 2017), were validated by qPCR analysis (Suppl. Fig. S5C).

As summarized in Fig. 7, while in HeLa cells the abundant KDELR2 is mainly involved in client retrieval, KDELR3 and KDELR1 exert additional regulatory roles, modulating AGR2 expression upon binding ERp44 and other unidentified ligands. Thus, a complex regulatory pathway centered on KDELRs can report on and adjust the composition of the ESP protein factory.

**Figure 7.**
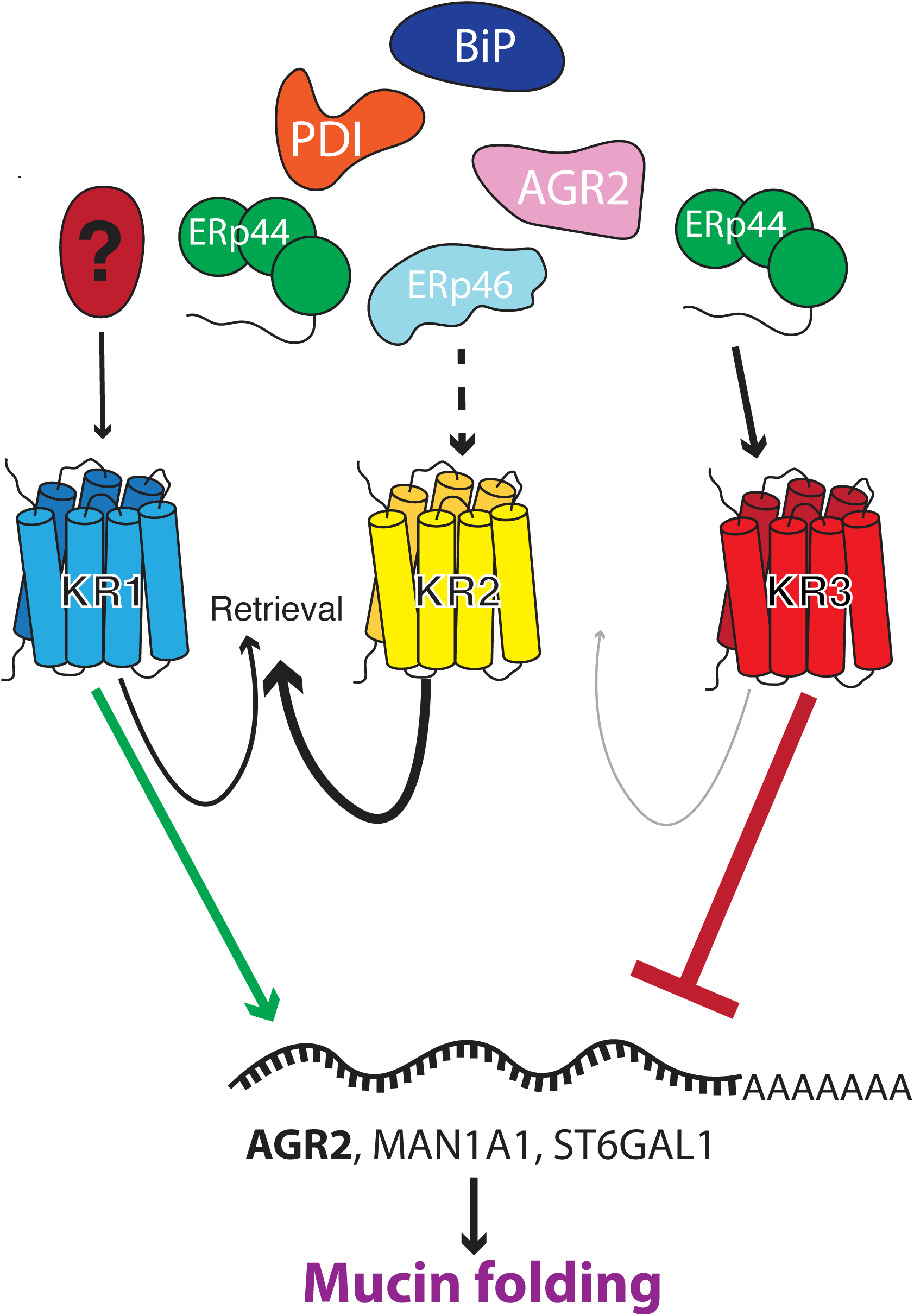
Functional specialization of human KDELRs. As summarized in the right part of the cartoon, ERp44 and KDELR3 act epistatically to inhibit AGR2 transcription. Instead, KDELR1 (on the right) can be stimulated by different chaperones to promote AGR2 transcription. In HeLa cells, the more abundant KDELR2 and KDELR1 sustain retrieval activity of diverse ER chaperones and enzymes.

## DISCUSSION

In metazoans, the diversification of the secretome imposed serious challenges for the ER protein factory. Numerous additional ubiquitous and cargo-specific folding assistants have evolved to ensure precision and efficiency in the production of ligands as well as receptors capable of transmitting the desired *inter-cellular* messages. Also, the prototypic yeast HDEL diversified, including KDEL and RDEL, while the KDELRs triplicated.

If preventing chaperone loss was the sole function of KDELR, why triplicate the efficient yeast module? Retrieval efficiency largely reflects the abundance of the human receptors, independent of their isoform. However, interactions between different KDELRs and their clients (each with a rather conserved tetrapeptide motif) can modify the ESP protein factory and its delivery compartment. Signals emanating from KDELR1 and KDELR3 promote and inhibit the accumulation of AGR2 (transcripts and proteins), respectively. ERp44, a protein regulated by pH and zinc during its cycling in the early secretory compartment (Amagai et al, 2023; Vavassori et al, 2013; Watanabe et al, 2019) is a privileged interactor of KDELR3, albeit not the sole. ERp44^KD^ phenocopies KDELR3^KD^ when AGR2 transcription is analyzed, suggesting that the two proteins serve epistatic functions in the emerging regulatory pathway. ERp44, but neither ERp46 nor Halo-RDEL, can rescue AGR2 transcriptional inhibition in HeLa^ERp44KO^ cells. Excitingly, rescue requires the presence of KDELR3. Thus, not all chaperones eventually retrieved by KDELRs dispatch the same signal(s). It will be of interest to dissect the binding kinetics and competition between KDELRs and chaperones and how these are influenced by the nature of their tetrapeptide motifs as well as by the binding of specific cargoes.

Our transcriptomic analyses confirm the special relationship between ERp44 and KDELR3. Large overlaps emerge when either component of the ERp44-KDELR3 axis is silenced. Considering the essential role of AGR2 in mucin biogenesis and Ire1β regulation (Cloots et al, 2024; Neidhardt et al, 2024), the regulatory circuitry we describe here may be relevant in inflammatory bowel disease, a condition in which AGR2 plays a crucial role (Al-Shaibi et al, 2021; Maurel et al, 2019; Zheng et al, 2006). Accordingly, genes related to O-glycosylation, an essential step in mucin biogenesis, are also significantly upregulated upon KDELR3^KD^ and ERp44^KD^.

Human KDELRs have long been considered as a single functional entity. However, recent studies suggested likely differentiation in their activities. Thus, the over-expression of KDELR1 and KDELR2, but not of KDELR3, stimulated extracellular matrix degradation (Ruggiero et al, 2015). KDELR2 and KDELR3, but not KDELR1, are targets of the Ire1α/Xbp1 pathway (Trychta et al, 2018). Mutations in KDELR1 and KDELR2 have been linked to lymphopenia (Siggs et al, 2015), and osteogenesis imperfecta, respectively (Efthymiou et al, 2021; van Dijk et al, 2020). More recently, KDELR2 was found upregulated in chronic obstructive pulmonary disease (Wu et al, 2024). Finally, mutations in KDELR3 are associated with type 2 diabetes (Altenhofen et al, 2023) and severe myopia (Yuan et al, 2024), and its loss, but not KDELR1 loss, impairs melanoma metastasis (Marie et al, 2020). Here we show that the three KDELRs are differentially involved in delivering signals from the ESP.

If the ER were a homogeneous pool of proteins retained through their C-terminal KDEL motifs, all should leak out of cells at similar rates upon inhibition of the three KDELRs. Remarkably, instead, ERp44 and ERp46 became barely detectable in cells devoid of all three KDELRs, whereas most BiP and PDI remained in the ER. We showed previously (Anelli et al, 2007; Palazzo et al, 2022; Tempio et al, 2021) that ERp44 exits the ER faster than other residents by binding to ERGIC53, a lectin that exerts rapid cycling by facilitating entry into COPII vesicles (Kappeler et al, 1997). BiP and other slow-moving residents tend to interact with one another as well as with cargoes in transit, likely forming larger condensates with slow mobility (Meunier et al, 2002; Reddy et al, 1996). Hence, the kinetics of recycling as well would impact KDELR-based signaling. Clients, pH, and/or zinc can modulate ERp44 recycling and, in turn, KDELR3 signals. A consequence of such differential recycling rates would be the generation of gradients along the ESP, depending on the balance between true retention in the ER (Meunier et al, 2002; Reddy et al, 1996) (formation of supramolecular complexes with slow diffusion), forward movement (bulk flow or assisted by cargo receptor) (Gomez-Navarro et al, 2020), and retrieval from the Golgi by KDELRs. Thus, not only the abundance of a chaperone and its cargo substrates but also the velocity at which it leaves the ER would modulate KDELR engagement. In the emerging scenario, the interactions between KDELRs, cycling chaperones and cargoes in transit may generate signals that allow adjusting the ESP composition to the proteostatic demand.

## METHODS

### Reagents

Unless otherwise specified, all reagents used were purchased from Sigma-Aldrich (St. Louis, Missouri, USA). Detailed information about the experimental protocols and the biotechnological reagents used, such as antibodies or oligonucleotides, are provided below in dedicated paragraphs.

### Cell lines and culture conditions

All the experiments were performed on human HeLa or Caco-2 cell lines originally supplied by ATCC. Genetic tests were performed to ensure their identity.

HeLa cells were cultured in Dulbecco’s Modified Eagle’s Medium (DMEM, Gibco, Invitrogen) supplemented with 10% Fetal Bovine Serum (FBS, Euroclone) and Penicillin/Streptavidin (Pen/Strep). Caco-2 cells were cultured in DMEM supplemented with 10% FBS and Gibco MEM Non-Essential Amino Acids (MEM NEAA, Thermo Fisher Scientific), without Pen/Strep.

For cell subculturing and plating, cells were rinsed with Dulbecco’s Phosphate Buffered Saline Solution (DPBS 1X, Corning) and detached with trypsin 0.05%-EDTA 0.53 mM (Corning). As a standard protocol, 1.5 mL were used for 10 cm plates, and proportionally lower volumes for smaller culture supports, incubating for 3 minutes at 37°C, 5% CO2 before inactivation with fresh culture medium.

### Plasmids

The plasmid encoding for SP-Halotag-RDRDEL was previously described ^(Mossuto^ ^et^ ^al, 2014)^. Halo-ERp44 was excised from pcDNA-Halo-ERp44 (Sannino et al, 2014) by NheI-EcoRV digestion, and inserted into pTRE2hyg (Clontech, Mountain View, USA). The resulting construct was verified by SmaI digestion and sequencing (Eurofins Discovery) before transfection. The construct obtained was used as a scaffold to perform Gibson assembly and substitute ERp44-RDEL with ERp46-KDEL (a kind gift from the late Professor Neil Bulleid).

For generating BioID2 tagged KDELRs, cDNAs of KDELR1, KDELR2, and KDELR3 were cloned from the cDNA library of HEK293T cells and subcloned into the pEF4/Myc-His vector B using the BamHI/EcoRI sites (Invitrogen). The BioID2 gene was PCR-amplified from the MCS-BioID2-HA plasmid (a gift from Dr. Kyle Roux; Addgene plasmid # 74224 ; http://n2t.net/addgene:74224 ; RRID:Addgene_74224) and inserted into the KDELR1-3 overexpressing plasmids using the KpnI/BamHI sites. Finally, a 13xGGGGS linker was inserted into the BglII/BamHI sites to generate the BioID2-13xGGGGS-KDELR1, 2, or 3 expressing plasmid.

To prepare the plasmids for overexpression of siRNA-resitant-KDELR1-3 (si-r-KDELR1-3), 4–6 bases (underlined) were alterted in the sequence of the pEF4/KDELR-MycHis plasmids by PCR-based mutagenesis as follows:

si-r-KDELR1 (CTA<u>T</u>CT<u>G</u>TA<u>C</u>AT<u>T</u>AC<u>G</u>AA<u>G</u>/CCAA<u>T</u>TA<u>T</u>AT<u>T</u>TC<u>T</u>CTCTA)

si-r-KDELR2 (AAAC<u>G</u>AT<u>A</u>CT<u>G</u>TA<u>T</u>TG<u>C</u>GA/CTA<u>T</u>CT<u>C</u>GC<u>G</u>TG<u>T</u>TC<u>G</u>TAT)

si-r-KDELR3 (GAA<u>C</u>TT<u>T</u>TA<u>C</u>GA<u>T</u>CA<u>G</u>ATT/AGAC<u>G</u>AT<u>C</u>AC<u>C</u>AC<u>A</u>CA<u>T</u>TA)

### Silencing assays

Cells were plated in 6 wells or 6 cm plates to have about 50-60% confluence on the day of transfection. Following the manufacturer’s instruction, cells were transfected using Lipofectamine RNAiMAX (Invitrogen). Duplexes for ERp44 downregulation have been previously described (Anelli et al, 2007); pre-designed duplexes were purchased from Thermo Fisher; for ERp46 TXNDC5 Silencer Select Pre-designed siRNA, s37649; for AGR2, s20692. The sequences of the duplexes used for silencing KDELRs are listed in Table 1. MISSION® siRNA Universal Negative Control (MERCK) was used as a control. 3 µL duplexes (20 µM) were used for each well of a 6-well plate. Cells were incubated overnight at 37% and 5% CO2. The morning after, the supernatant was replaced with 2 mL of fresh medium. Cells were harvested and lysed 72 hours after silencing. Some batches of certified HeLa cells (i.e. HeLa Kyoto) tended to be less viable upon KDELR silencing.

**Table 1:**
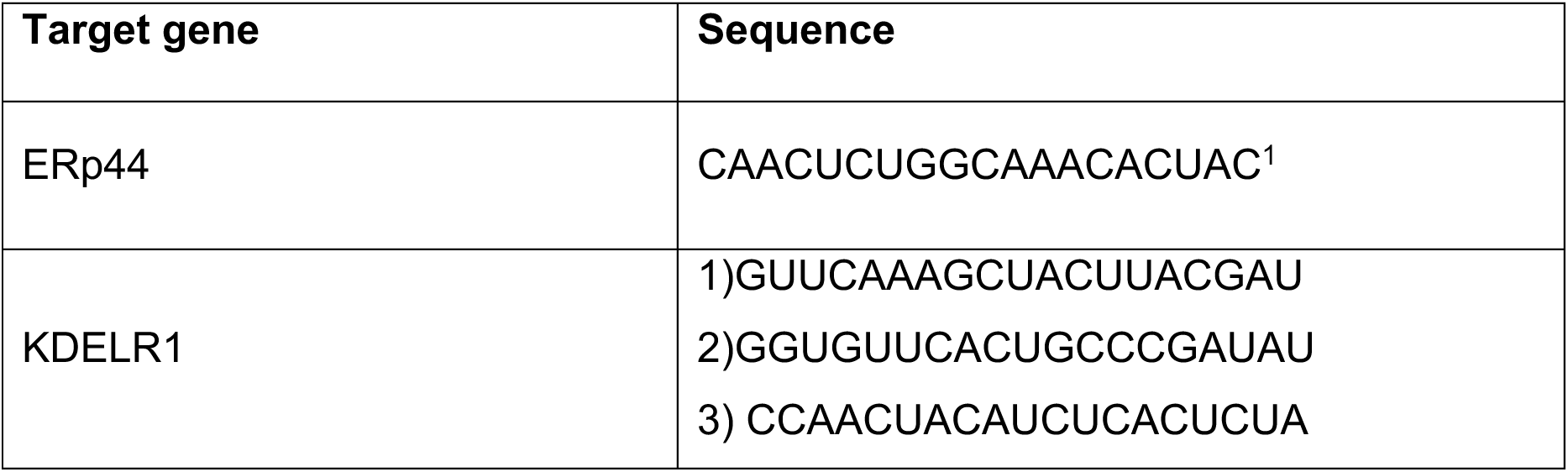
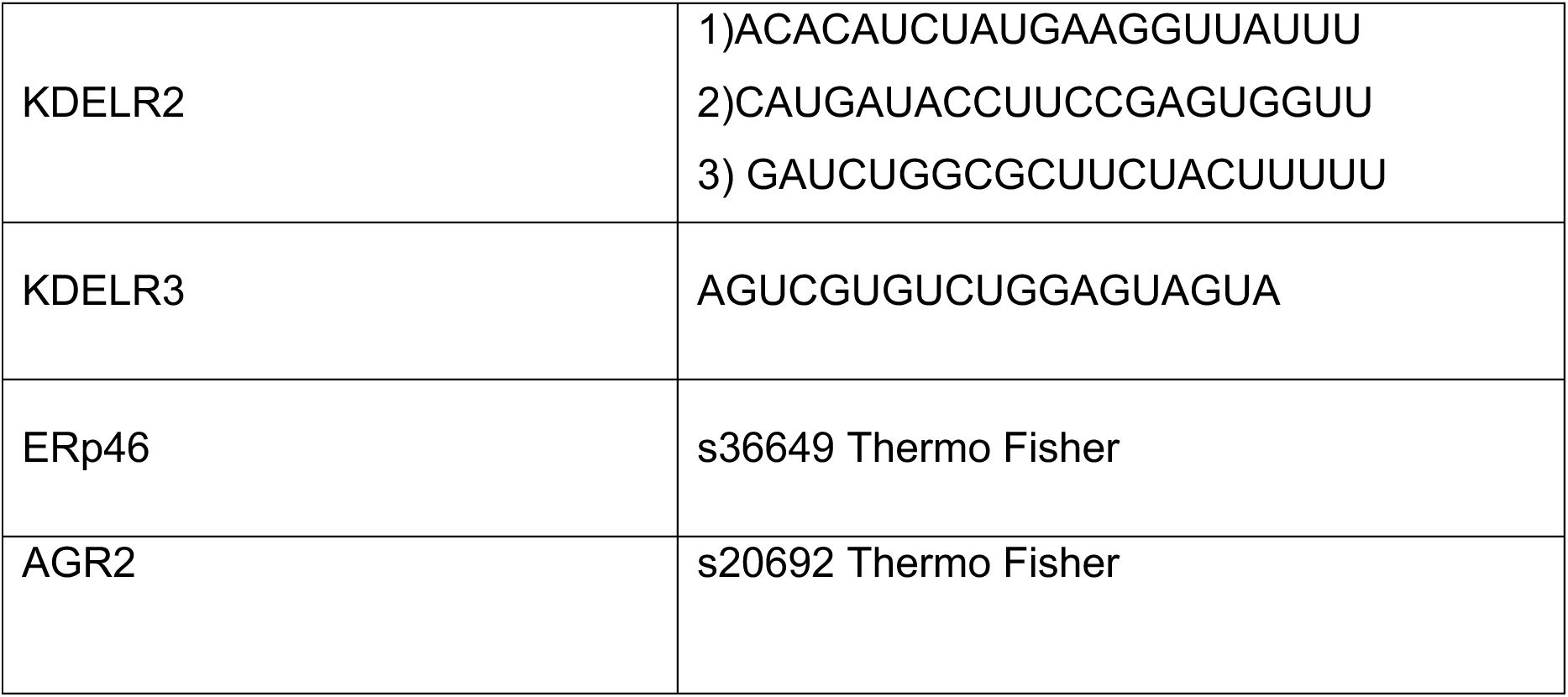
Oligos (silencing assay)

### Knockdown/rescue experiment for KDELRs-Myc

Two types of siRNA duplex (#1 and #2) were simultaneously transfected to knock down each KDELR isoform. Silencer Select negative control No. 2 siRNA (Cat No. 4390846, Thermo Fisher Scientific) served as a control (siControl).

HeLa Kyoto cells (0.4×10^5^ cells) were plated onto a coverslip in a well of a 6-well plate. After incubation for 18 h, cells were transfected with the si-r-KDELRs-Myc plasmids or an empty vector using FuGENE HD (Promega) and incubated for an additional 6 h. Cells were further transfected with 5 nM of siRNAs using Lipofectamine RNAiMAX. The cells were then incubated for 36 h and fixed with 4% PFA/PBS for 10 min.

### BioID2 experiments

HEK293T cells (2.0×10^6^ cells) were plated onto 10-cm dishes. After incubation for 24 h, cells were transfected with 5 μg of plasmid overexpressing BioID2-KDELR1, 2, or 3. Cells were further incubated for 16 h, and the culture media was exchanged with fresh DMEM containing 2 μM of biotin. After incubation for 16 h, cells were washed with PBS three times, harvested, and lysed in 1 mL of lysis buffer containing 50 mM tris(hydroxymethyl)aminomethane (Tris), pH 7.4, 500 mM NaCl, 0.4% sodium dodecyl sulfate (SDS), 1 mM dithiothreitol (DTT), and 1× protease inhibitor cocktail. Cell extracts were then sonicated for 1 min using an Astrason 3000 instrument (Misonix Incorporated) at output level 3.0, followed by the addition of TritonX-100 to a final concentration of 2%. After 1 min sonication at output level 3.0, an equal volume of 50 mM Tris (pH 7.4) was added, and samples were sonicated for a further 30 s. Supernatants were collected in 15 mL tubes after centrifugation at 22,000 × g for 15 min and rotated with 15 μL of streptavidin agarose beads overnight at 4°C. Beads were collected by centrifugation at 3,500 × g for 3 min and washed twice with 1 mL of wash buffer containing 2% SDS for 8 min, once with 1 mL of wash buffer containing 0.1% sodium deoxycholate, 1% Triton X-100, 500 mM NaCl, 1 mM ethylenediaminetetraacetic acid (EDTA), and 50 mM 4-(2-hydroxyethyl)-1-piperazineethanesulfonic acid (pH 7.5), and twice with wash buffer 3 containing 50 mM Tris (pH 7.4) and 50 mM NaCl. Beads were treated with 1× Laemmli buffer containing 2 mM biotin, and the resulting eluates were denatured at 98°C for 5 min. Proteins collected using streptavidin agarose beads from four 10-cm plates of HEK293T cells were resolved by SDS-PAGE on 12.5% gradient precast gels and detected by silver staining using a Silver stain II kit. The gel was then cut into six pieces per lane, and gel pieces were digested into peptides using trypsin. The resultant peptide fragments were then separated by high-performance liquid chromatography (HPLC) on an Advance UHPLC instrument (Bruker). The molecular masses of peptides were determined by mass spectrometry (MS) using an Orbitrap Velos Pro instrument (Thermo Fisher Scientific). Proteins with emPAI values >0.03 were identified as plausible interactors of each KDELR isoform. Proteins were considered positive if more than three digested peptides were identified, and hence their subcellular localizations and functions were further investigated using the UniProt database (https://www.uniprot.org/).

### Secretion and cell lysis assays

Secretion assays were performed by removing the cell culture medium, rinsing the plate with DPBS 1X, and adding either 1 or 2 mL of Opti-MEM in each well of a 6-well plate or a 6 cm plate, respectively. Cells were incubated for 4 hours at 37°C in 5% CO2. Supernatants (SN) were then collected, additioned with 10 mM N-Ethylmaleimide (NEM) and cOmplete^TM^ Protease Inhibitor Cocktail (Roche), and centrifuged to remove cell debris. Samples were precipitated using trichloroacetic acid to both concentrate proteins and remove possible contaminants (salts and detergents) before loading in SDS-PAGE gels.

Cells were collected and lysed in RIPA buffer (50 mM Tris-HCl pH 7.5, 150 mM NaCl, 1% NP-40, 0,5% Deoxycholate, 0,1% SDS) supplemented with 10 mM NEM and cOmplete^TM^ Protease Inhibitor Cocktail. Post-nuclear supernatants were collected for further analysis.

Cell lysates were quantified following a standard Bicinchoninic Acid (BCA) Protein assay, using purified Bovine Serum Albumin (BSA) for the calibration curve.

### Western Blot Assays

Protein samples were loaded onto pre-cast Invitrogen SDS-PAGE 4-12% or 4-15% gradient gels. Routinely, 40 µg of cell extracts/lane were loaded, and resolved under reducing conditions. After electrophoresis, proteins were transferred onto nitrocellulose membranes, either via a wet transfer in an electric field, 300 mA for 1 h and 30 minutes at 4°C using a buffer containing 20% methanol and 10% standard Tris-Glycine buffer, or via a dry transfer using the Trans-Blot Turbo Transfer System by Bio-Rad setting the mixed molecular weight transfer program. After transfer, the nitrocellulose membrane was saturated with 5% milk in DPBS 0.1% Tween and then incubated with the primary antibodies (Table 2). After careful washes, appropriate secondary antibodies Alexa Fluor^®^ (Invitrogen) conjugated with diverse fluorophores (647, 546 nm) or chemoluminescence antibodies for UVITEC (Southern Biotech, anti-mouse HRP, anti-rabbit HRP) were used. The acquisition was performed at the correct wavelength using Typhoon FLA9000 Biomolecular Imager (GE Lifesciences) or, in the case of HRP antibodies, the emitted light was acquired with UVITEC imager (Uvitec). When required by the experimental design, densitometric analysis was performed with ImageJ software, using tubulin bands for lane normalization.

**Table 2:** Antibodies (immunoblotting)

| Antibody | Species | Dilution |
| --- | --- | --- |
| ERp44 | Mouse, clone 36C9 (in-house, Anelli 2007) | 1:500 in PBS 0.1% Tween |
| ERp46 | Rabbit (Sigma) | 1:500 in PBS 0.1% Tween |
| PDI | Rabbit (Braakman lab) | 1:1000 in PBS 0.1% Tween |
| AGR2 | Rabbit (proteintech) | 1:500 in PBS 0.1% Tween |
| GAPDH | Mouse (SantaCruz) | 1:1000 in PBS 0.1% Tween |
| KDEL | Mouse (Enzo) | 1:500 in PBS 0.1% Tween |

### Immunofluorescence and Pearson’s correlation coefficient analysis

HeLa Kyoto cells (0.17×10^5^) were seeded onto a 12-well plate with glass cover slips over the well. Cells transfected with siRNAs targeting KDELRs for 24 h were fixed by 4% paraformaldehyde (PFA) for 15 min, permeabilized by 0.1% Triton X-100/PBS solution for 30 min, and blocked by 2% FCS/PBS solution for 1 h at room temperature. Subsequently, cells were incubated with antibodies against ERp44 and GM130 or calnexin diluted in HIKARI A solution overnight at 4°C. CF568-conjugated anti-mouse IgG and CF488-conjugated anti-rabbit IgG antibodies (Biotium) were used as secondary antibodies. Nuclei were stained with 4’,6-diamidino-2-phenylindole (DAPI) diluted in 2% FCS/PBS at room temperature for 10 min. Stained cells were embedded in Fluoro-KEEPER antifade reagent before observation. Cell images shown in Figure 2 were obtained using an FV-1000 confocal laser scanning microscope (OLYMPUS) equipped with a UPLSAPO 60× silicon-oil immersion objective lens (NA 1.3) and processed with ImageJ software. To optimize the images, microscopy settings were adjusted in each single experiment. Pearson’s correlation coefficients of ERp44 and GM130 signals were calculated by JACoP (Just Another Colocalization Plugin).

For the knockdown/rescue experiments, fixed cells were permeabilized with PBS containing 0.1% TritonX-100 for 15 min at RT and blocked with 2% FBS/PBS for 1 h at RT. Cells were then incubated with anti-Myc polyclonal antibodies (MBL; cat# 562, 1:5000 diluted in 2% FBS/PBS) for 2 h at RT. Cells were then washed with PBS three times and further incubated with an anti-ERp44 monoclonal antibody (2D5, 1:50 diluted in HIKARI A) overnight at 4°C. After washing cells three times with PBS, cells were incubated with CF488-labeled anti-mouse IgG and CF568-labeled anti-rabbit IgG antibodies for 1 h at RT. DNAs were labeled with DAPI. Coverslips were mounted onto slide glasses using Fluoro-KEEPER antifade reagent. Fluorescence images were obtained with a laser scanning confocal microscope (LSM980, Carl Zeiss) equipped with a Plan Apochromat 40x lens (NA = 0.95). Signal intensities of ERp44 in each cell were quantified using Fiji (NIH).

### PCR and Real-Time Quantitative PCR

Total RNA was extracted with 1 mL of phenol-based reagent TriFast (Euroclone, Cat#EMR507100), lysing cells directly on the culture plate, and samples were then stored at –80°C. RNA was separated in the aqueous phase by the addition of chloroform to the samples and centrifugation. The collected SN were precipitated with 200 μL isopropanol and centrifuged. The pellets were then washed with 1 mL of 75% ethanol before resuspension in RNAse-free water (usually 20 μL, more for bigger RNA pellets).

Retro-transcription to cDNA was performed with M-MLV Reverse Transcriptase and random primers (Promega, Cat#M1701; Cat#C1181) following the manufacturer’s protocol for 2 μg RNA.

Real-time semi-quantitative PCR was performed with iTaq Universal SYBR Green Supermix (Bio-Rad) and with couples of gene-specific primers (specifics are in Table 3). Each reaction was performed in duplicate in a total volume of 10 μL. Amplification was performed with the BioRad CFX96^TM^ System. The amplification was performed with a denaturation step of 10 min at 95°C, followed by 40 cycles of PCR amplification (15 s at 95°C, 1 min at 60°C), and by a step-by-step temperature increase from 60°C to 95°C, to calculate the melting curve (Dalla Torre et al, 2024).

**Table 3:**
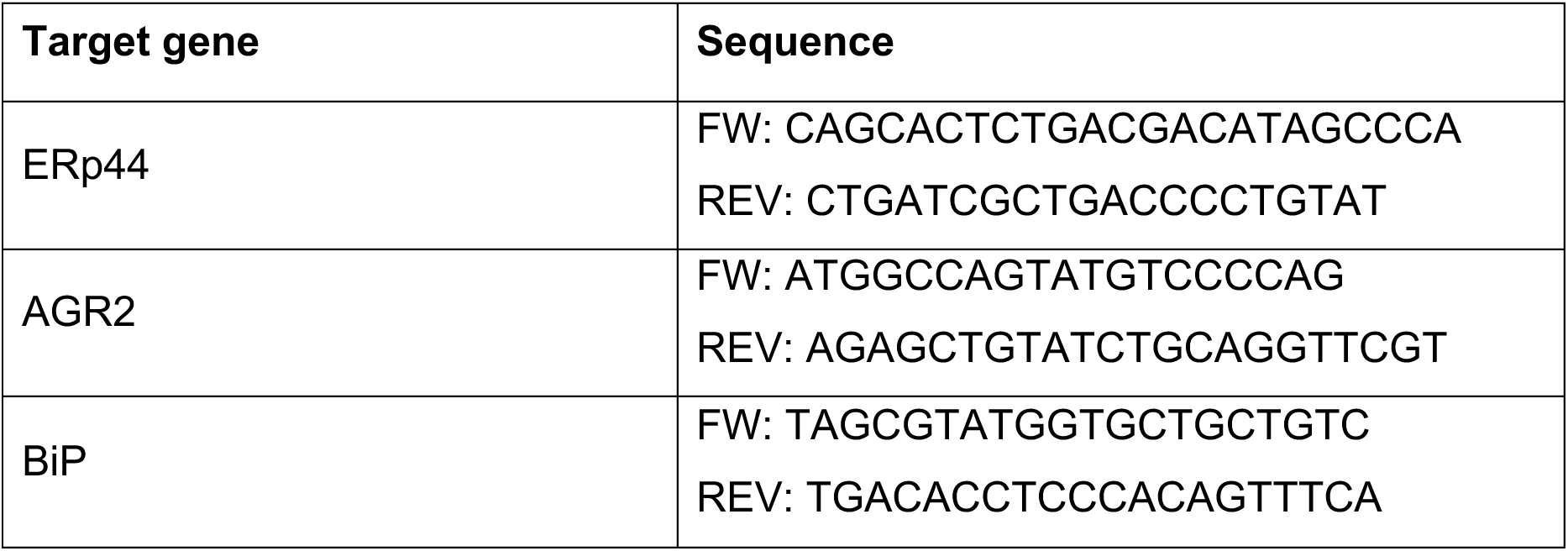
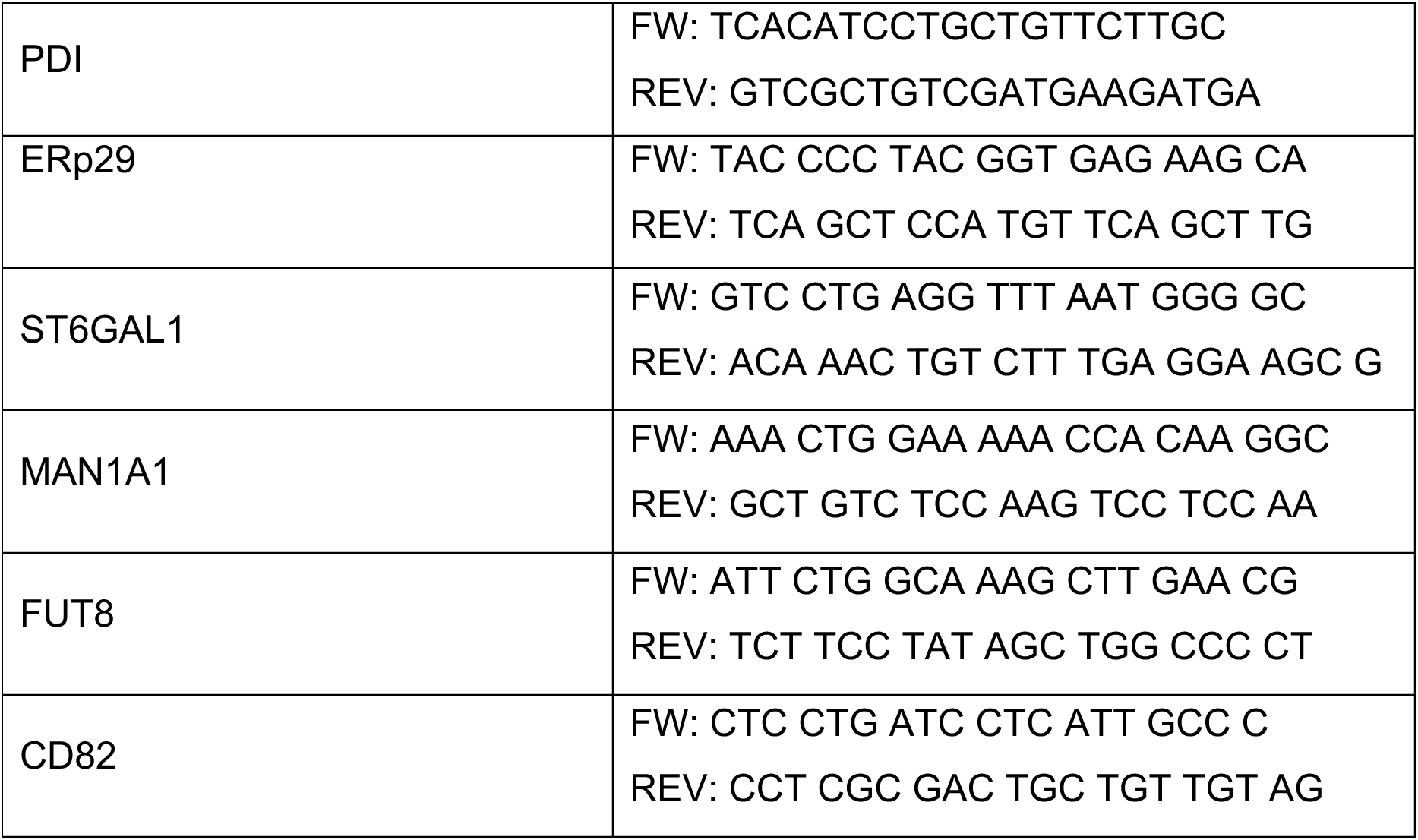
Real-time qPCR primers.

GAPDH was chosen as a reference gene to perform relative quantifications. For all experiment conditions, the ΔCt was calculated by subtracting the Ct of the reference gene from the Ct of the gene of interest. The values obtained were then used to calculate the ΔΔCt by subtracting the ΔCt of the control condition from the other conditions, with the results of each gene representing the net relative increase/decrease relative to the control. These numbers were finally used to calculate the fold gene expression value (2^-ΔΔCt^) often used in literature. Values obtained in the different experiments were averaged and plotted.

qRT-PCR primers used for KDELRs were obtained from Qiagen (Hilden, Germany): Hs_KDELR1_1_SG QuantiTect Primer Assay (QT00090811) for KDELR1, Hs_KDELR2_1_SG QuantiTect Primer Assay (QT00092715) for KDELR2, Hs_KDELR3_1_SG QuantiTect Primer Assay (QT00097790) for KDELR3. All the other primes were purchased from Metabion (Planegg, Germany) and are listed in Table 3.

### Transcriptomics

HeLa cells were subjected to silencing as described above. Five replicates for each condition were used. 72 hours after silencing cells were collected and RNA extracted using the RNeasy Plus Kits for RNA Isolation (QIAGEN, ID: 74134) and quantified by Nanodrop.

Libraries of cDNA were prepared with TruSeq mRNA Library Prep Kit (Illumina) and sequenced using the Novaseq instrument (Illumina) 100 bp single-end up to 30 million reads. After trimming the adapter sequences from reads (Trimmomatic), sequences were aligned to the human hg38 genome using the STAR (v2.5.3a) (https://github.com/alexdobin/STAR/) and counted using *feature Counts* (http://bioconductor.org/packages/release/bioc/html/Rsubread.html) with the gene annotation from Gencode (version31). The R/Bioconduction package ‘limma’ was used to conduct differential expression analysis using standard procedures (Ritchie et al, 2015).

Raw sequencing data and raw counts are available on the Gene Expression Omnibus repository, accession number: GSE289834 (token to review GEO accession GSE289834: wlcbuiqsxpijzun).

Functional GO Enrichment was performed using the R/Bioconductor package ClusterProfiler (Xu et al, 2024) and heatmap plots were made with ‘pheatmap’ package (Kolde, 2018).

Enrichment analysis of the genes in overlap in Figure 6C was made using the Enrichr online server (Xie et al, 2021). Gene Ontology Cellular Compartments used in Figure S4 are listed in Table 4.

**Table 4:** GO compartments.

| Compartment | Gene Ontology Cellular Compartments |
| --- | --- |
| **Golgi** | cis-**Golgi** Network (GO:0005801)<br>cis-**Golgi** Network Membrane (GO:0033106)<br>Clathrin Coat Of trans-**Golgi** Network Vesicle (GO:0030130)<br>trans-**Golgi** Network (GO:0005802)<br>trans-**Golgi** Network Membrane (GO:0032588)<br>trans-**Golgi** Network Transport Vesicle (GO:0030140)<br>**Golgi** Cis Cisterna (GO:0000137)<br>**Golgi** Cisterna (GO:0031985)<br>**Golgi** Cisterna Membrane (GO:0032580)<br>**Golgi** Lumen (GO:0005796)<br>**Golgi** Medial Cisterna (GO:0005797)<br>**Golgi** Membrane (GO:0000139)<br>**Golgi** Stack (GO:0005795)<br>**Golgi** Trans Cisterna (GO:0000138)<br>**Golgi**-associated Vesicle (GO:0005798)<br>**Golgi**-associated Vesicle Membrane (GO:0030660) |
| **ERGIC** | Endoplasmic reticulum-**Golgi** Intermediate Compartment Membrane (GO:0033116)<br>COPII-coated ER To **Golgi** Transport Vesicle (GO:0030134)<br>ER To **Golgi** Transport Vesicle Membrane (GO:0012507) |
| ER | **Endoplasmic Reticulum** (GO:0005783)<br>Rough **Endoplasmic Reticulum** (GO:0005791)<br>Rough **Endoplasmic Reticulum** Membrane (GO:0030867)<br>Signal Recognition Particle, **Endoplasmic Reticulum** Targeting (GO:0005786)<br>Smooth **Endoplasmic Reticulum** (GO:0005790)<br>Cortical **Endoplasmic Reticulum** (GO:0032541)<br>Cytoplasmic Side Of **Endoplasmic Reticulum** Membrane (GO:0098554)<br>**Endoplasmic Reticulum** Lumen (GO:0005788)<br>**Endoplasmic Reticulum** Membrane (GO:0005789)<br>**Endoplasmic Reticulum** Tubular Network (GO:0071782)<br>Extrinsic Component Of **Endoplasmic Reticulum** Membrane (GO:0042406)<br>Luminal Side Of **Endoplasmic Reticulum** Membrane (GO:0098553) |
| Glycosylation | Protein C-linked **Glycosylation** Via 2'-alpha-mannosyl-L-tryptophan (GO:0018406)<br>Protein C-linked **Glycosylation** Via Tryptophan (GO:0018317)<br>Protein N-linked **Glycosylation** (GO:0006487)<br>Protein N-linked **Glycosylation** Via Asparagine (GO:0018279)<br>Protein O-linked **Glycosylation** (GO:0006493)<br>Protein O-linked **Glycosylation** Via Serine (GO:0018242)<br>Protein O-linked **Glycosylation** Via Threonine (GO:0018243)<br>Regulation Of Protein **Glycosylation** (GO:0060049)<br>Protein **Glycosylation** (GO:0006486) |

## ACKNOWLEDGMENTS

We thank Anush Bakunts, Johannes Buchner, Emma Fenech, Chiara Giannone, Chiara Gramegna Tota, Alberto Luini, Luca Rampoldi, Céline Schaeffer, Maya Schuldiner, and Elisa Speranza for helpful discussions and suggestions, Claudio Fagioli and Andrea Orsi for technical help. This work was partly supported by grants from AIRC (IG 23285) and MIUR/PRIN (XA5J5N) to RS, AMED-CREST (21gm1410006h0001) to KI, PNRR-M 4, C 2, I 1.1 funded by NextGenerationEU under PRIN2022MZJR9X to MS. This work was partly supported also by the Cooperative Research Project Program of the Medical Institute of Bioregulation, Kyushu University.

## AUTHOR CONTRIBUTIONS

FCP, KI, RS, and TA conceived and designed the project; FCP, YA, MDT, XH, and TT performed most of the wet experiments and analyzed the data critically; JGM and MM performed and interpreted the transcriptomics and proteomics analyses respectively; MS and MF provided important conceptual contributions. FCP, YA, KI, RS, and TA wrote the manuscript. All authors discussed the results and commented on the manuscript. TA and RS supervised the project.

The authors declare no conflict of interest.

## DATA AVAILABIITY

The datasets and computer code produced in this study are available in the following database:

- Gene Expression Omnibus repository, accession number: GSE289834 (token to review GEO accession GSE289834: wlcbuiqsxpijzun).

**Table S1 KDEL-like motif-bearing proteins identified by BioID2 technology**

KDEL-like motif-bearing proteins identified by BioID2 technology are listed. In case more than three peptides are detected, and emPAI values are >0.03 by mass spectrometry, the proteins are considered to be positive in association with KDELRs. The identified proteins are classified according to the specificity of the KDELR isoforms and are shown in different colors (orange, common to KDELR1, 2, and 3; green, common to KDELR1 and 2; cyan, common to KDELR1 and 3; purple, common to KDELR2 and 3). Subcellular localizations and functions are assessed based on the UniProt database (https://www.uniprot.org/). Selected KDEL-like motif-bearing proteins are also shown in Figure S3.

**Table S2 Gene Ontology Biological Process Functional Enrichment.**

GO BP overrepresentation analysis of Differentially Expressed Genes (FDR < 0.05 and log2FC > 1) with ClusterProfiler in the silencing samples.

---

**Figure S1.**
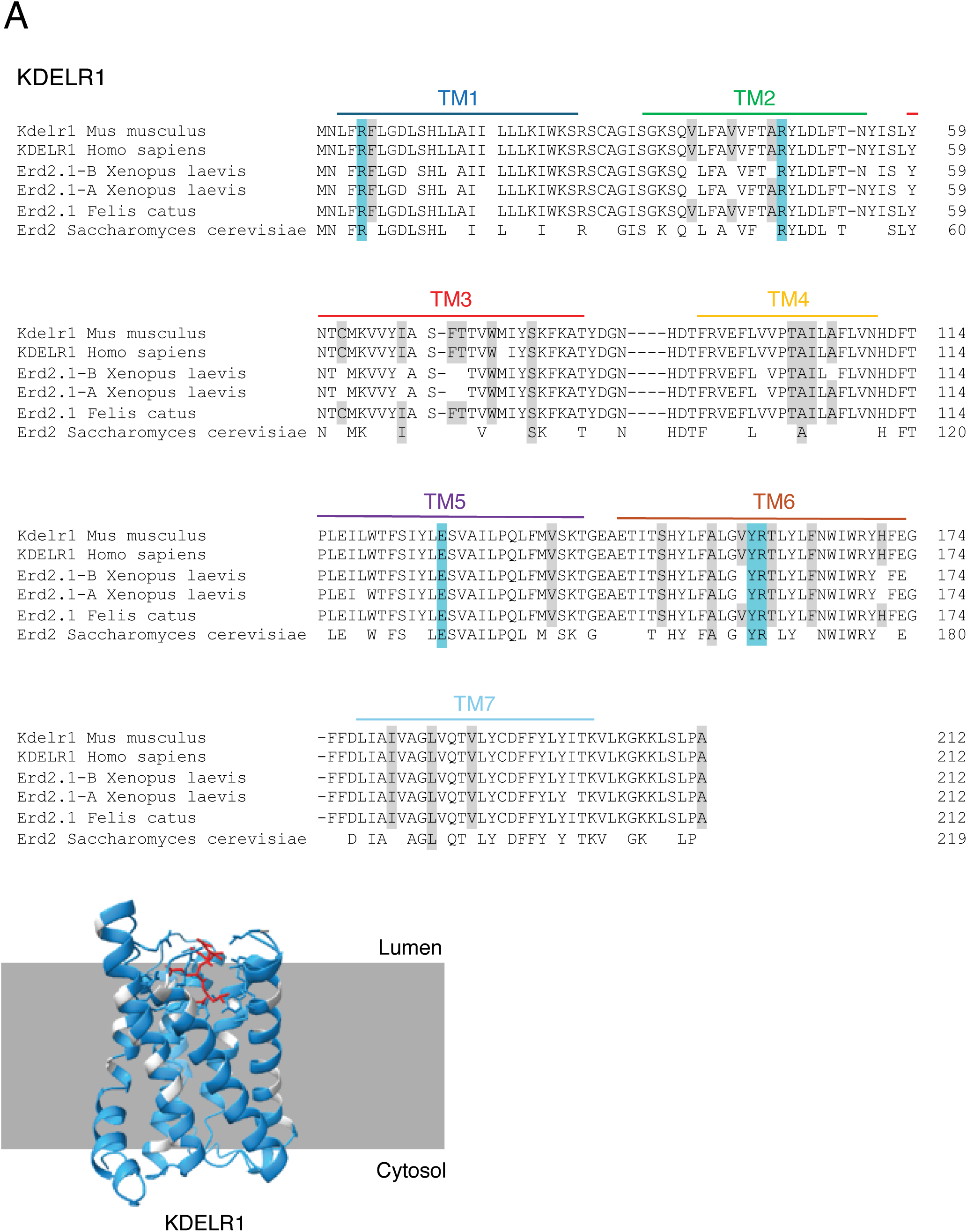
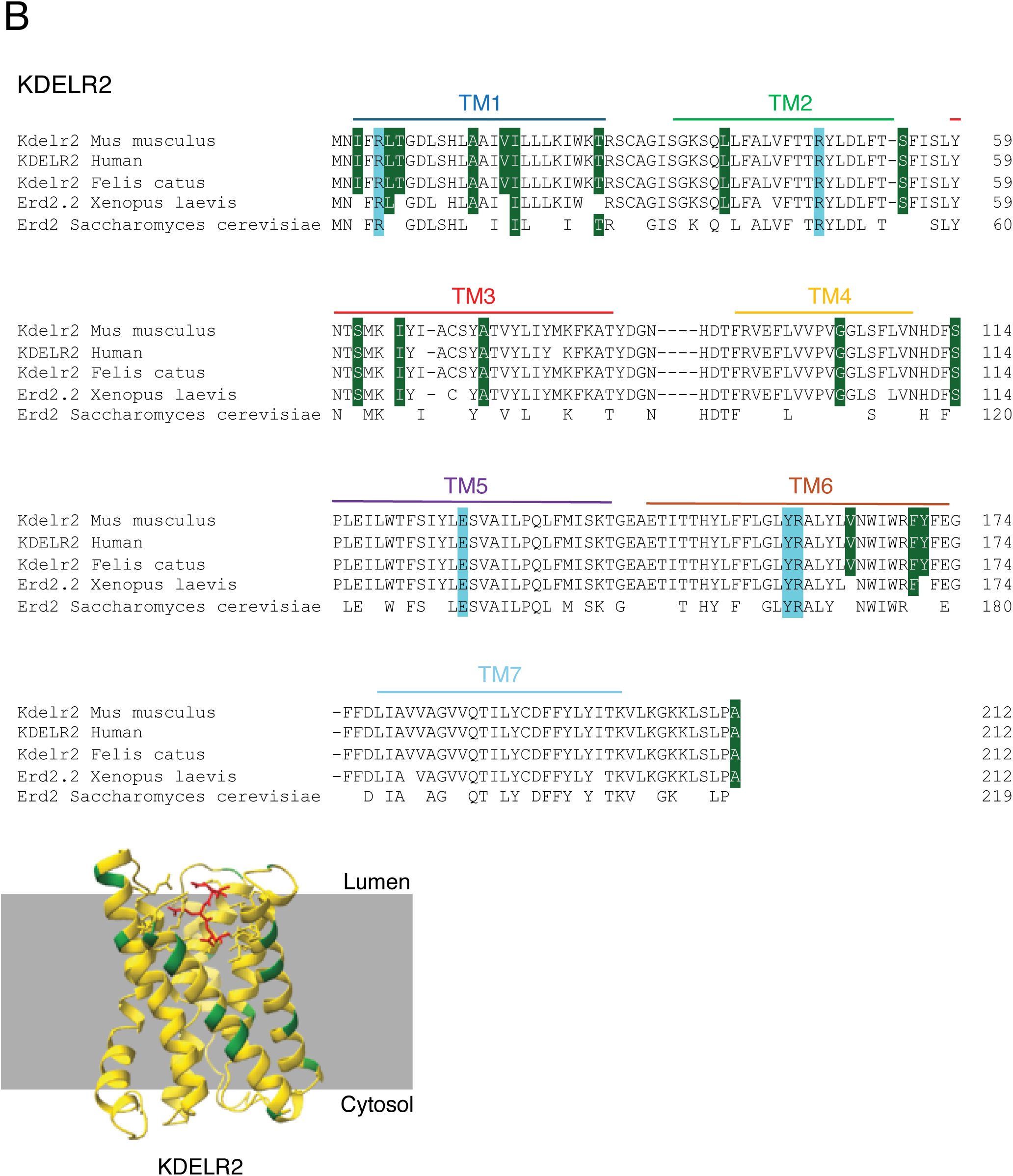
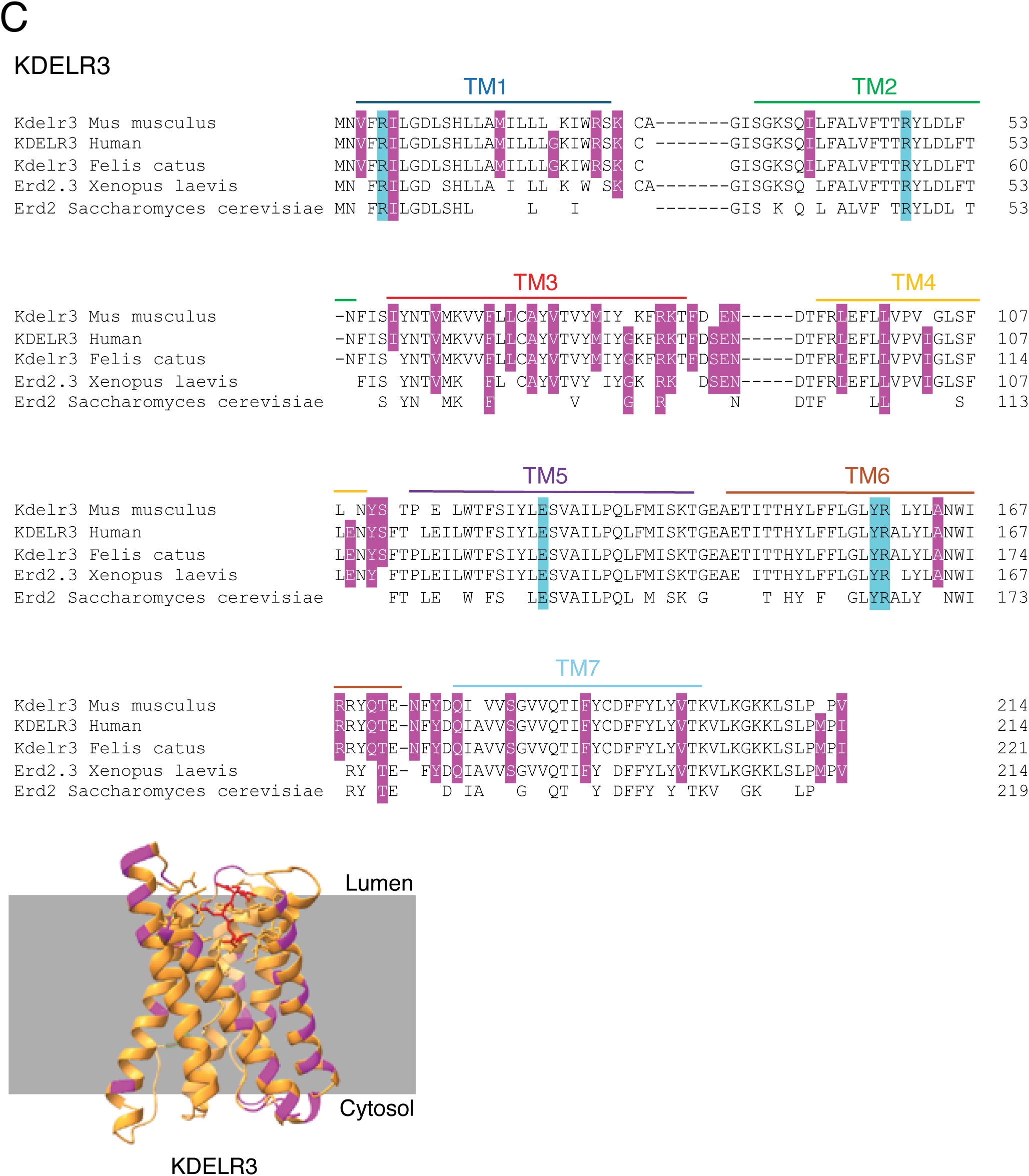
Phylogenetically conserved differences hallmark the three KDELRs. A) *Sequence alignment of the amino acidic sequence of KDELR1 in different species*. Upper part: the amino acidic sequences of KDELR1 in different species (*Mus musculus, Homo sapiens, Xenopus laevis, Felis catus, and Saccharomyces cerevisiae ERD2*) were aligned with *Geneious Prime*. The conserved differences, the residues involved in the KDEL binding and the TMs are highlighted by the same color code used in Fig. 1. Bottom part: the three-dimensional structure of the human KDELR1 is predicted using the AI tool Alphafold 3. Its features are explained in Fig. 1. B) *Sequence alignment of the amino acidic sequence of KDELR2 in different species*. For KDELR2, the conserved differences are highlighted in green. Additional color code is as described for panel A and Fig. 1. C) *Sequence alignment of the amino acidic sequence of KDELR3 in different species*. For KDELR3, the conserved differences are highlighted in purple. Additional color code as in panels A and B and Fig. 1.

**Figure S2.**
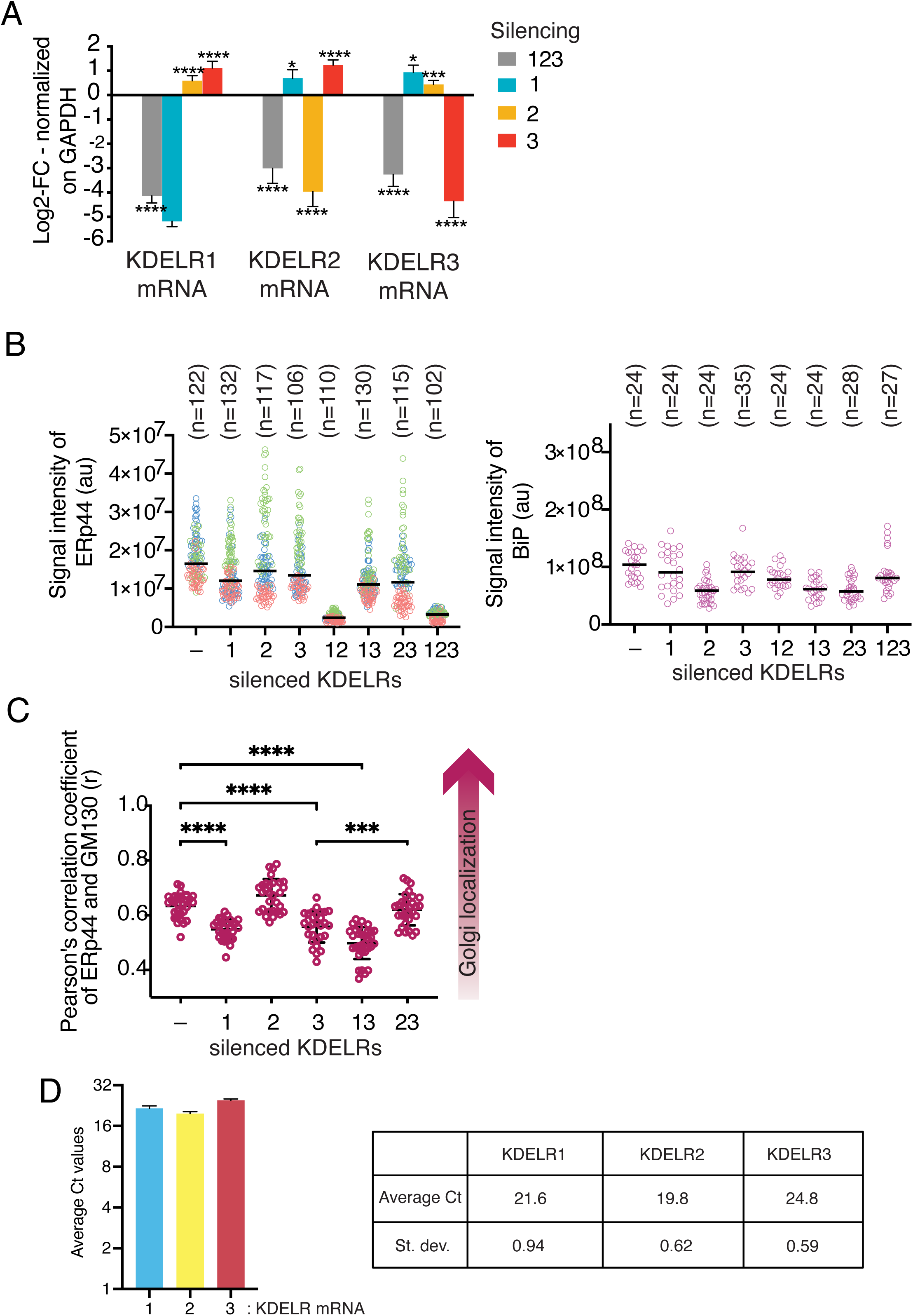
Different expression levels and functions of human KDELRs. A) *Knockdown efficiency of siRNA targeting of each KDELR isoform.* The abundance of mRNAs of each KDELR isoform was quantified by RT-qPCR, and normalized relative to untreated controls. Results are means ± SD of at least three independent experiments. Statistical significance was calculated by one-way ANOVA followed by Dunnett’s multiple comparison test (ns, not significant; \**p* = 0.0391; \*\*\**p* = 0.0002; \*\*\*\**p* = 0.0001). B) *Changes in the localization of ERp44 and BiP upon isoform-specific KDELR silencing*. The signal intensities of endogenous ERp44 (left) and BiP (right) under various KDELR^KD^ conditions were quantified based on the immunofluorescence signals shown in Figures 2C and D. C) *ERp44 proceeds along the secretory pathway upon KDELR silencing.* Pearson’s correlation coefficients of endogenous ERp44 and GM130 under various KDELR^KD^ conditions were calculated based on the immunofluorescence signals shown in Figure 2C. The ERp44 signals in double KDELR12^KD^ and KDELR123^KD^ cells were too weak to unequivocally localize and quantify. Statistical significance was calculated by one-way ANOVA followed by Tukey’s multiple comparison test (\*\*\*\**p* <0.0001). The experiment was repeated three times independently, and the results were consistent each time. D) *Different basal levels of the three KDELRs in HeLa cells.* The average Ct and SD of 10 different RT-qPCR experiments were calculated and shown both as a table and column chart. Clearly, KDELR2 is more abundant than KDELRs 1 and 3.

**Figure S3.**
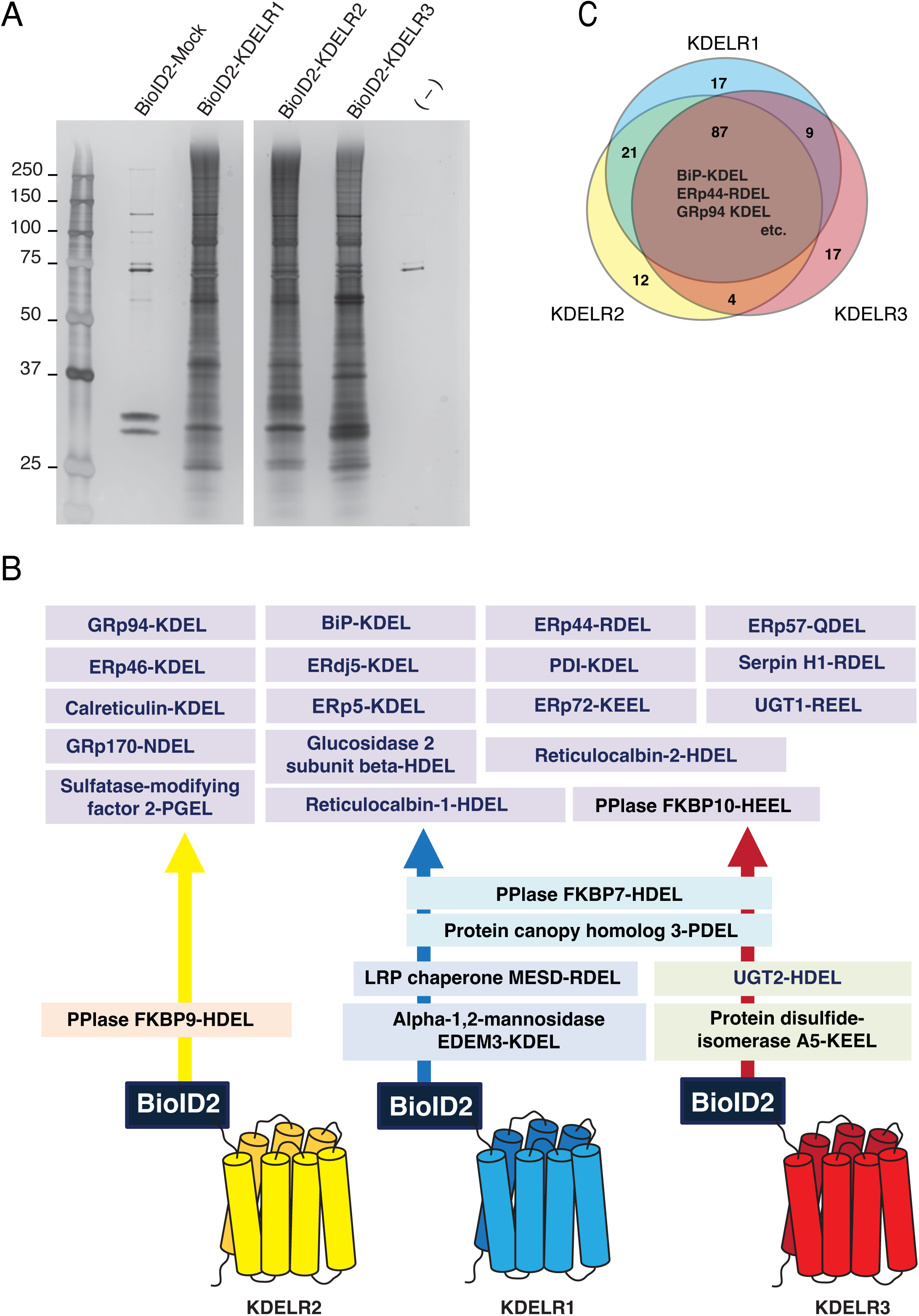
Proximity networks of KDELR isoforms 1, 2 or 3 identified by BioID2 technology. A) Biotinylated proteins recovered from HEK293T cell lysates overexpressing each of BioID2-KDELR1/2/3 were separated by SDS-PAGE and visualized by silver staining. Gel slices were subjected to trypsin digestion and mass spectrometry analysis for the identification of interacting proteins. B) KDEL-like motif-bearing proteins identified by the above method are classified based on their preferential binding to the different KDELR isoforms, as shown in different colored rectangles. C) The image summarizes the number of KDEL-like motif-bearing proteins associated to each KDELR isoforms. See also Table S1, which shows a list of proteins identified in the BioID2 experiment shown in panel A.

**Figure S4.**
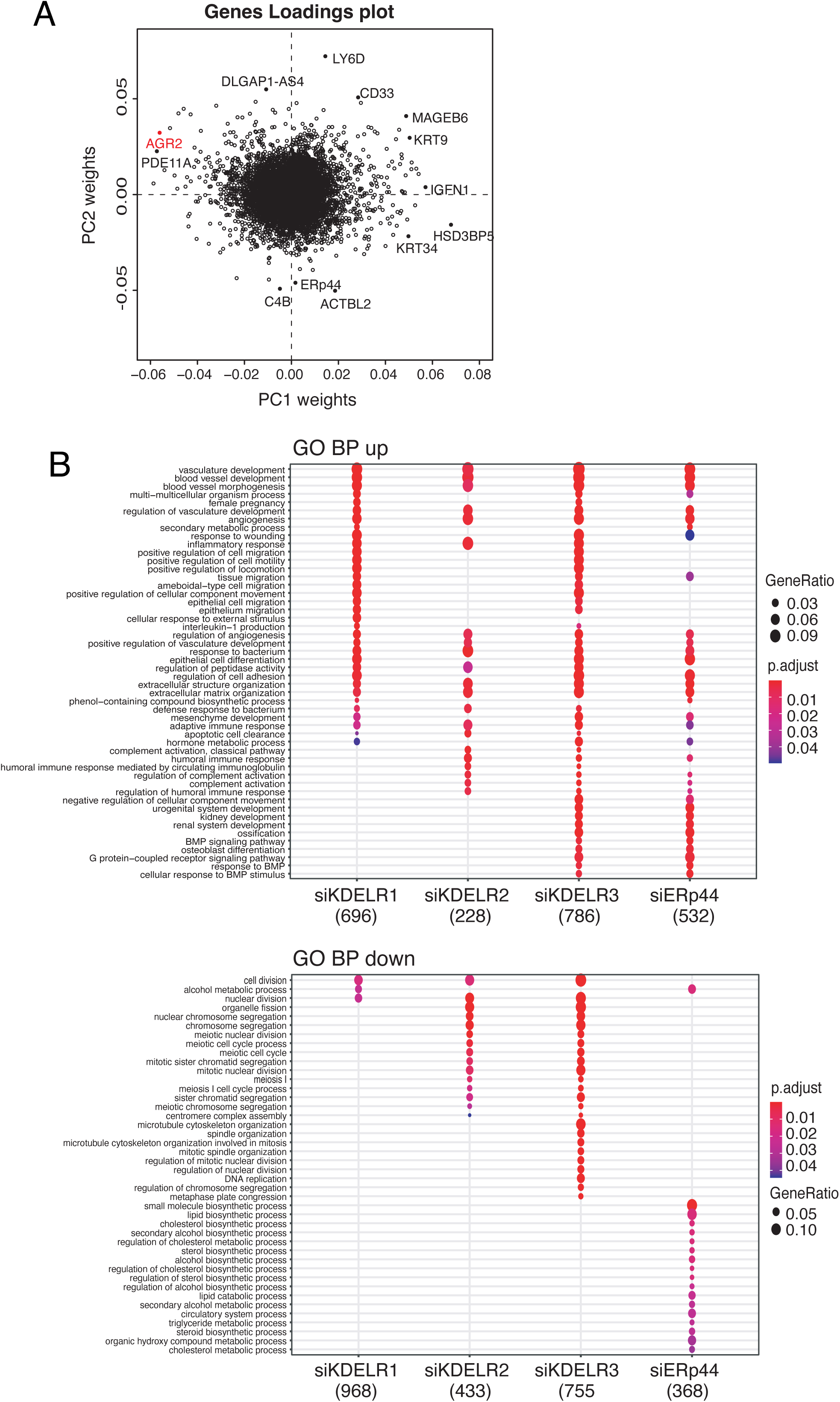
Transcriptional differences after KDELRs or ERp44 downregulation. A) The image highlights the genes whose differential expression governs the PCA variance in Figure 6A. B) Gene Ontology Biological Process Enrichment. Dotplot of top 20 (ordered by FDR) significant biological processes in UP (top panel) and DOWN (bottom panel) regulated genes (FDR < 0.05 and log2FC >1). The panel highlights mainly processes upregulated both upon KDELR3^KD^ and ERp44^KD^ as well as processes downreguated both upon KDELR2^KD^ and KDELR3^KD^.

**Figure S5.**
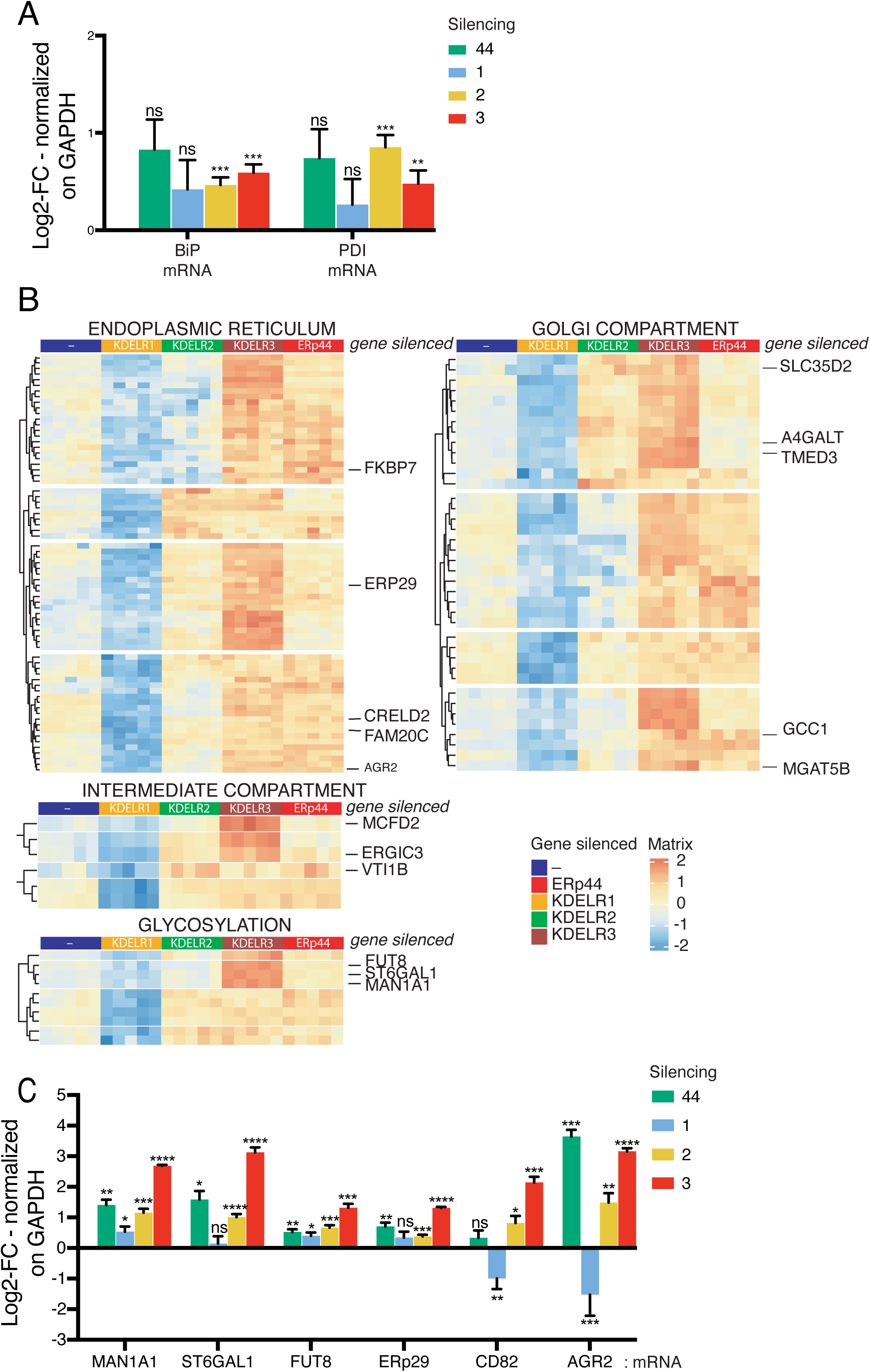
Transcriptional changes induced upon ERp44 or isoform-specific KDELR^KD^. The abundance of transcripts originating from the indicated genes was quantified by qRT-PCR experiments with specific oligos. A) BiP and PDI show a mild increase in all conditions, possibly reflecting an unspecific response to gene silencing. B) Heatmap of expression levels (log2 Normalized, Z-score scaled) of genes behaving as AGR2 (upregulated in KDELR3^KD^ and ERp44^KD^ and downregulated in KDELR1^KD^ with a FDR ≤0.05) belonging to the indicated lists of Gene Ontology Cellular Compartments, (ER, Golgi, Intermediate compartments and glycosylation). See Methods for further details. C) KDELR3^KD^ significantly induced the accumulation of MAN1A1, ST6GAL1, FUT8 (glycosylation), ERp29, and -as expected-AGR2 (chaperoning). CD82/KAI1, a tetraspannin involved in tumor progression ^(Marie^ ^et^ ^al, 2020; Zhu^ ^et^ ^al, 2017)^, was significantly downregulated upon KDELR1^KD^.

